# Repurposing without Reinventing the Wheel - Ensemble Models for Differential Analysis

**DOI:** 10.1101/2025.04.07.647549

**Authors:** Suvo Chatterjee, Erina Paul, Ziyu Liu, Jialin Gao, Piyali Basak, Arinjita Bhattacharyya, Chitrak Banerjee, Himel Mallick

## Abstract

Inspired by ensemble models in machine learning, we propose a general framework for aggregating multiple distinct base models to enhance the power of published differential association analysis (DAA) methods. We demonstrate this approach by augmenting popular DAA models with one or more biologically motivated alternatives. This creates an ensemble that bypasses the challenge of selecting an optimal model and instead combines the strengths of complementary statistical models to achieve superior performance. Our proposed ensemble learning approach is platform-agnostic and can augment any existing DAA method, providing a general and flexible framework for various downstream modeling tasks across domains and data types. We performed extensive benchmarking across both simulated and experimental datasets spanning single-cell gene expression, bulk transcriptomics, and microbiome metagenomics, where the ensemble strategy vastly outperformed non-ensemble methods, identified more differential patterns than the competing methods, and displayed good control of false positive and false discovery rates across diversified scenarios. In addition to highlighting a substantial performance boost for state-of-the-art DAA methods, this work has practical implications for mitigating the so-called reproducibility crisis in omics data science. An open-source R package implementing the ensemble strategy is publicly available at https://github.com/himelmallick/DAssemble.

## 1 Introduction

A longstanding goal of any statistical analysis of omics data is the identification of molecular or biochemical features associated with phenotypes, exposures, health outcomes, and other important covariates. These associations are typically investigated using differential association analysis methods, which entail identifying features that display differential expression or abundance patterns across samples. The terminology varies by omics data type: for example, differential expression analysis applies to transcriptomics, differential abundance analysis to microbiome studies, and analogous methods exist for proteomics and metabolomics. Other variants, such as differential distributional analysis^14^ or differential variability analysis ^47,66^, are also common. Without loss of generality, we refer to this fundamental problem as differential association analysis (DAA) and the associated differential features as differentially associated (DA) features.

In its simplest form, DAA compares the per-feature omics measurements of a given population between two experimental groups, such as cases versus controls or healthy versus diseased samples. The statistical foundation of DAA is two-sample hypothesis testing, a well-studied problem for over 100 years ^91^. Yet, performing DAA reliably has proven particularly challenging due to the statistical properties of high-throughput omics datasets, which are often characterized by high dimensionality, count and compositional data structures, sparsity, overdispersion, and hierarchical, spatial, and temporal dependencies, among others^28,29^. Indicative of these challenges, numerous DAA methods developed in recent years yield remarkably different results. For instance, while one method may detect hundreds of DA features in a particular dataset, another may detect none. Until now, no consensus has emerged on the best-performing DAA method^76^—a surprisingly consistent observation across omics domains, including microbiome ^31^, gene expression^29,50^, and spatial omics^13^, among others.

Although identifying DA features may seem straightforward for a given dataset, the variability in results across different DAA methods raises important questions about the reliability of published approaches. Notably, benchmarking studies frequently provide conflicting recommendations, making it challenging for practitioners to select the most appropriate method ^22,26,28,29,106^. Numerous base models, ranging from parametric to nonparametric, exist to capture multifaceted biological data properties and experimental design considerations, further complicating the choice for practitioners. In practice, it is generally impossible to know a priori which base model will perform best for a given problem. Inspired by ensemble models in machine learning ^96^, we hypothesize that a weighted combination of multiple candidate DAA models into an ensemble method can mitigate these issues. Theoretically, such an ‘ensemble learner’ is expected to perform as well as or better than any individual candidate model ^96^, thereby enhancing the performance of existing differential analysis methods.

In addition to boosting performance, ensemble models can also enhance interpretability, particularly when combining two drastically different yet synergistic models that capture distinct properties of the data, greatly expanding the scope of biological hypotheses that can be tested. For example, in single-cell differential expression, binarized expression profiles often provide a more robust representation of biological variation compared to count measurements^7,21^—a phenomenon observed across many other omics fields, including microbiome and spatial transcriptomics, where appropriate logistic regression models are commonly used for differential analysis after data binarization^41,76^. Similar to ensemble learning and Bayesian model averaging ^20,33^, aggregated models can help address uncertainty in model selection, potentially mitigating the so-called ‘reproducibility crisis’ in omics data science, where studies often fail to replicate and methods show inconsistency in achieving consensus ^26,28,31,50,76^.

Recently, a few studies have taken initial steps to construct consensus models for various down-stream analysis tasks, including normalization^89^, microbiome differential abundance ^70^, single-cell differential expression^5,49^, cell-type deconvolution^19^, multi-omics integration^30^, whole genome sequencing (WGS) association studies^55^, and foundation models^69^, among others. One approach, known as differential distributional analysis ^14,48^, provides a unified framework that simultaneously assesses differential expression, variability, and other important aspects of the data. Another set of methods leverages ensemble learning concepts to integrate multiple DAA approaches (typically more than two), combining their cooperative strengths through holistic integration ^5,49^. While the former often faces computational bottlenecks and limited flexibility, the latter suffers from an unclear statistical rationale for combining models, resulting in ‘black box’ ensembles that are difficult to interpret. We argue that the careful selection of candidate models is crucial for capturing the non-redundant aspects of broad-ranging modeling paradigms and ensuring downstream interpretability, particularly in DAA, where prediction is seldom the primary goal, unlike in machine learning. Moreover, scalability can be severely compromised when multiple models are combined indiscriminately without a statistical basis ^10^, highlighting the need for a nuanced selection of candidate models based on statistical and biological complementarity.

In this paper, we present a flexible, statistically motivated ensemble modeling strategy for differential analysis that aggregates diverse base models, leveraging their strengths without compromising performance, interpretability, or scalability. Our proposed ensemble learning approach is platform-agnostic and can enhance any existing DAA method, providing a versatile and adaptable framework for a range of downstream modeling tasks across platforms and data types. Experimental results demonstrate that our ensemble strategy significantly improves statistical power while maintaining Type I error and false discovery rates (FDRs) at nominal levels across multiple modalities, including bulk RNA sequencing (RNA-Seq), single-cell RNA-Seq, and metagenomics, providing a unified and general framework for differential analysis of multiplatform omics data. An open-source R package implementing the ensemble strategy is publicly available at https://github.com/himelmallick/DAssemble.

## 2 Results

### 2.1 Ensemble models augment published methods by integrating complementary statistical models

There is an overwhelmingly large literature and algorithms already available on differential analysis based on different modeling techniques and cultures^65,76,84^. However, standard DAA methods for inferring differential expression or abundance often suffer from the uncertainty of one single model. Our primary goal in this article is not to add one more new methodology to the existing toolbox but instead to develop an ensemble strategy that utilizes the mutually informative statistical properties of published methods. Such ensembles are key in various estimation approaches, including machine learning and Bayesian statistics. Unlike machine learning, where prediction is the primary goal, our focus here is on improving the performance of published DAA methods while also enhancing the interpretability of differential association results.

As most DAA methods are based on generalized linear models (GLMs), our approach seamlessly integrates with standard pipelines across domains. Instead of relying on a single model, we consider an ensemble of candidate models following standard preprocessing steps (**Fig. 1**). For evidence synthesis, we incorporate various pooling methods from the meta-analysis literature. Unlike standard pipelines, the ensemble approach provides both omnibus significance and model-specific significance, enriching differential analysis results with multivariate effect sizes (**Fig. 1**). In this paper, we focus on building two-part ensembles (**Table 1**); however, the approach is adaptable to any number of distinct models (**Fig. 1**). Similarly, while we use per-feature p-values for ensemble aggregation (**Methods**), the methodology is generalizable to any inferential procedure, including alternatives such as Bayes factors. Specifically, we apply three p-value aggregation methods: (1) Cauchy Combination ^56^ (CC), (2) Minimum P-value Combination^36^ (MC), and (3) Stouffer Combination^90^ (SC), and we correct for multiple hypothesis testing using FDR correction ^3,4^ (**Methods**). As a cautionary note, we included the SC method as a representative from the meta-analysis literature^42^. However, SC was originally developed for independent tests—an assumption clearly violated in the ensemble framework. For practical purposes, we recommend using other dependence-aware combination methods, such as the CC method or its variants^44,81,101^.

**Figure 1:**
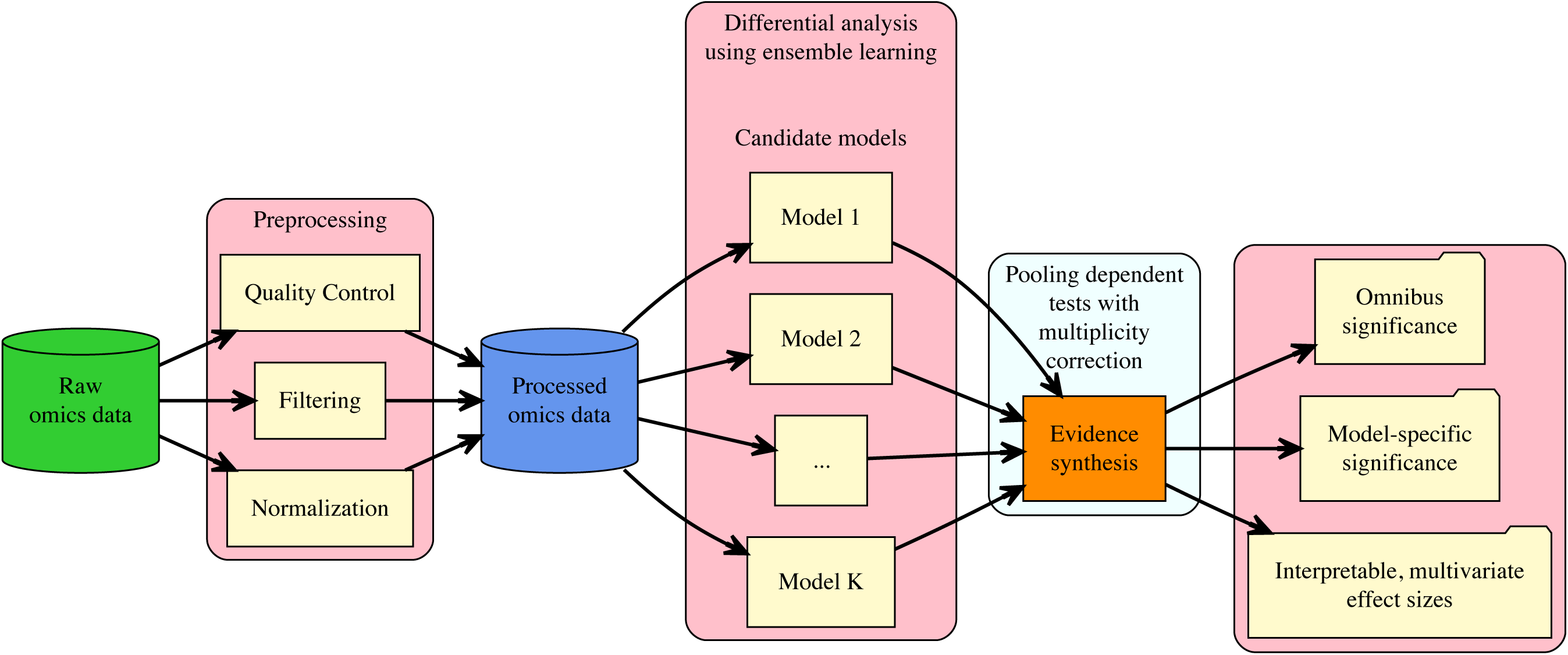
Ensemble models for differential association analysis. Starting with quality-controlled omics data, an ensemble model sequentially fits multiple, carefully curated candidate models. Individual model-specific effect sizes are collected, and statistical evidence is pooled using various combination methods, followed by multiplicity correction. In addition to providing both model-specific and omnibus significance results, ensemble models enable interpretable, multivariate effect sizes for further downstream analysis.

**Table 1:**
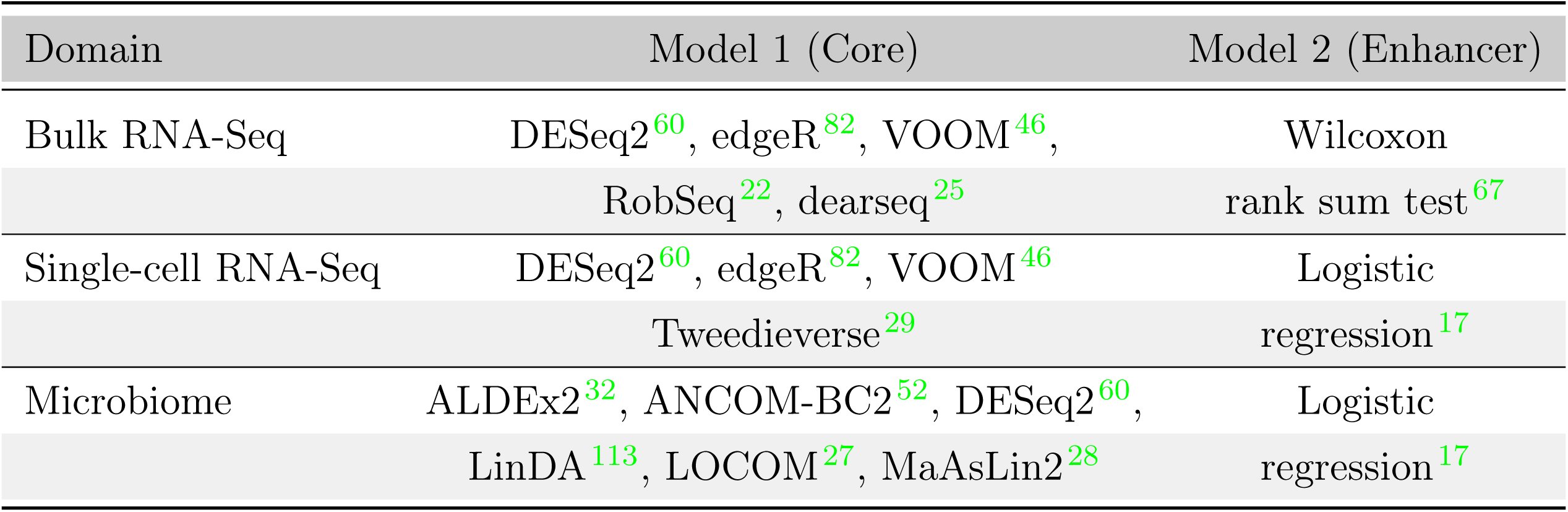
A summary of two-part ensembles considered in this study. Overview of the base models, enhancer models, and the three p-value combination strategies—Cauchy Combination (CC), Stouffer’s Combination (SC), and minP Combination (MC)—used to construct the ensemble variants.

To avoid creating a ‘black box’ while ensuring that results remain both robust and interpretable, we recommend selecting models based on their statistical and biological complementarity, as combining similar models may provide only incremental value. For example, a two-part ensemble using two different count-based GLMs, such as Poisson and Negative Binomial, may yield limited additional insights beyond a modest performance improvement. In contrast, integrating a count-based GLM with logistic regression can provide a more comprehensive view of the data, offering a multivariate effect size that captures both the magnitude of differential expression and differential detection^35^.

To illustrate how a well-curated ensemble can enhance the performance of differential association by effectively integrating compatible base models, we showcase various use cases in diversified scenarios (**Table 1**). In the context of single-cell differential expression (DE), we use a two-part ensemble with logistic regression for binarized gene expression profiles^7,21^, integrated with popular GLMs that model gene expression counts, motivated by the compelling performance of logistic regression in recent single-cell benchmarks^7,21^. For bulk RNA-Seq, we combine various parametric models with the Wilcoxon rank sum test^67^, motivated by the excellent performance demonstrated by this nonparametric method in recent studies^22,50^. Finally, for microbiome differential abundance detection, we supplement published methods with a logistic regression model for prevalence (presence-absence) modeling, driven by its strong empirical performance and the ecological relevance of presence-absence patterns in microbial communities^64,70^.

Notably, one of the models in our two-part formulation can be considered the core model, while the other serves as the enhancer (**Table 1**). As the name implies, this concept is inspired by enhancers in genetics, which are short DNA sequences capable of binding specific proteins (transcription factors) to increase the likelihood of a nearby gene being transcribed, thereby acting as regulatory elements to boost gene expression ^72^. Similar to enhancers in genetics, which can be located either upstream or downstream of the target gene and function independently of their orientation on the DNA strand, the selection of core and enhancer components in our ensemble formulation is likewise arbitrary.

### 2.2 Two-part ensemble models maintain FDR while enhancing power in bulk RNA-Seq differential expression analysis

To evaluate the effectiveness of the ensemble strategy in detecting differentially expressed genes (DEGs) in bulk RNA-Seq data, we assessed five core models used for DE analysis (**Methods**; **Table 1**). Two of these models, DESeq2^60^ and edgeR ^82^, are based on negative binomial (NB) GLMs. Another core model, dearseq ^25^, operates under a nonparametric framework, while the remaining two, VOOM ^46^ and Robseq ^22^, are based on linear models. Our ensemble strategy for DE analysis in bulk RNA-Seq data combines these core models with the classical Wilcoxon rank sum test^100^ (WLX). WLX was selected as an enhancer due to recent evidence demonstrating its strong FDR control and robust performance in DE analysis ^50,51^. This combination resulted in three ensemble variants for each core model: CC, SC, and MC. For example, for edgeR, these variants are edgeR + WLX (CC), edgeR + WLX (SC), and edgeR + WLX (MC), with each variant reflecting a different p-value combination method used to determine gene-specific significance.

To evaluate these models across cross-context settings, we generated synthetic bulk RNA-Seq data using the nonparametric simulator SimSeq ^2^. SimSeq utilizes an existing bulk RNA-Seq dataset as a template to generate synthetic data that captures key features of the original dataset, including mean-variance trends and both technical and biological variability (**Methods**). Each simulated dataset consisted of 10, 000 genes, with 10% designated as DE genes. We assessed model performance at two sample sizes, *N* = 40 and *N* = 100, equally divided into two experimental groups. The Cancer Genome Atlas (TCGA) Colorectal Cancer (CRCA) dataset (**Methods**) was used as a template for SimSeq to generate the synthetic data. To enhance result robustness and minimize random sampling errors, each simulation scenario was replicated 100 times. After generating the synthetic data, we evaluated two key aspects of model performance: (i) the ability of the core models and their ensemble variants to detect DE genes (sensitivity/power) and (ii) their ability to control the FDR at the nominal 5% level (**Methods**).

Several noteworthy observations emerged. For a small sample size (*N* = 40), we found that the statistical power of most core models and their CC and MC ensemble variants was similar, indicating that the CC and MC variants did not significantly enhance sensitivity (Figure 2A). In contrast, the SC variant of all ensemble models showed a notable increase in sensitivity, with power improvements of 15% to 20%. Similarly, for a larger sample size (*N* = 100), the SC variant of each ensemble model yielded the greatest improvement in statistical power compared to other variants (Figure 2C). These observations support our central hypothesis that the ensemble approach effectively enhances sensitivity across both small and large sample sizes, achieving up to a 20% increase in power.

**Figure 2:**
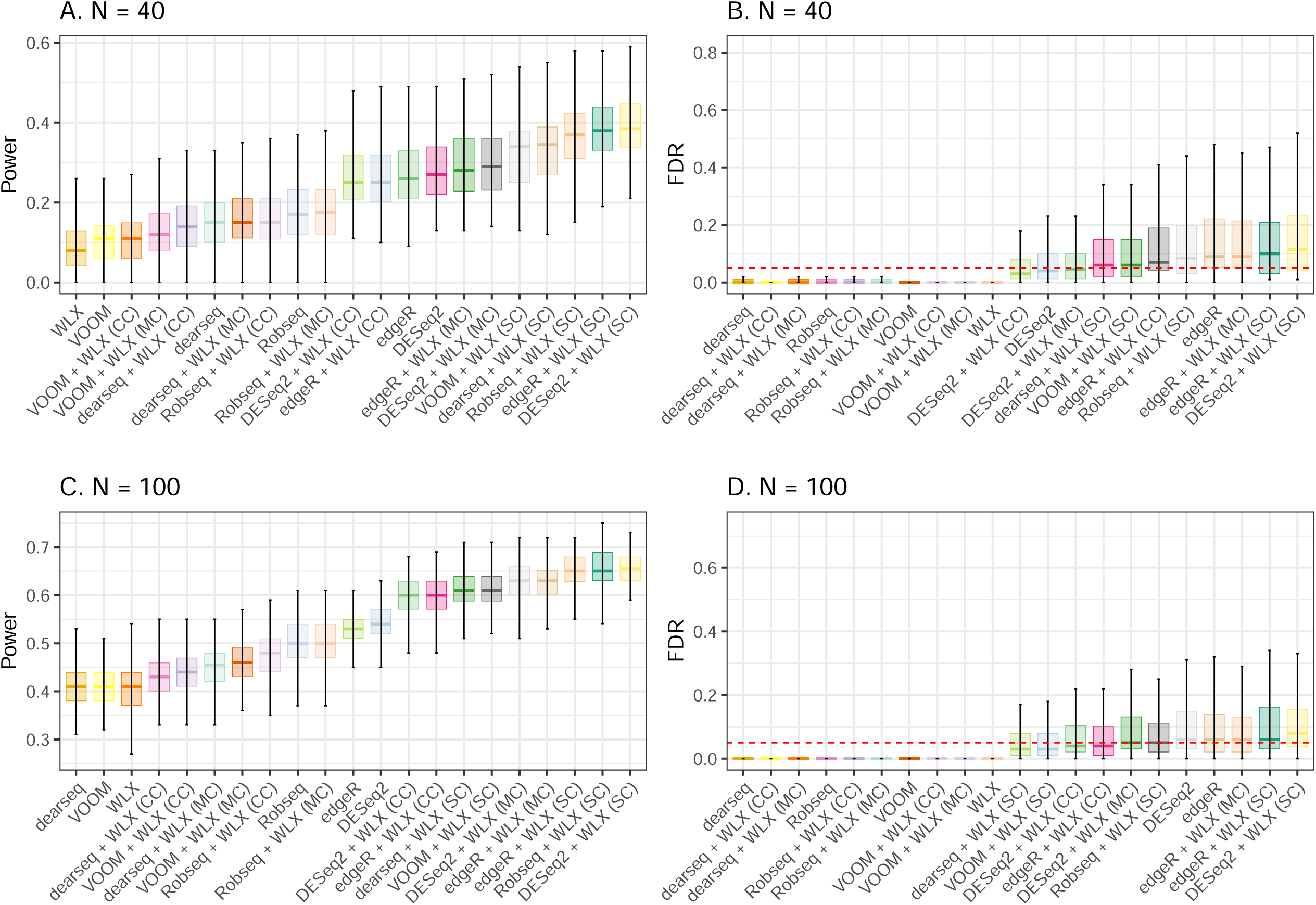
Ensemble models boost statistical power while controlling FDR in bulk RNA-Seq DE analysis. We simulated 100 synthetic datasets under two different sample sizes, with the true proportion of DE genes set to 10%. Panels **A** and **C** show boxplots of statistical power from 20 DE methods, ordered by increasing median sensitivity for sample sizes *N* = 40 and *N* = 100, respectively. Panels **B** and **D** show the corresponding FDR. Methods on the right side of the power boxplots generally demonstrate higher sensitivity. Methods maintaining FDR below the nominal 0.05 threshold appear below the red dotted line.

In terms of false discovery, all core models controlled the FDR at the nominal 5% threshold when *n* = 40, except for edgeR, which exhibited an inflated FDR of 0.09 (Figure 2B). Similarly, the CC and MC variants of all ensemble models controlled the FDR at the nominal level, except for the CC and MC variants of edgeR + WLX. In contrast, the SC variants of the ensemble models—dearseq + WLX, VOOM + WLX, and Robseq + WLX—showed marginal FDR inflation, with median FDRs between 0.06 and 0.08, while the SC variants of DESeq2 + WLX and edgeR + WLX were moderately liberal, with median FDRs between 0.08 and 0.11. Under larger sample sizes (*N* = 100), we observed that all core models and their SC, MC, and CC ensemble variants controlled the FDR at the target 5% level, except for edgeR + WLX and DESeq2 + WLX (Figure 2D). Consistent with the literature^50^, NB-based models and their ensemble variants (edgeR, DESeq2, edgeR + WLX (MC), edgeR + WLX (SC), and DESeq2 + WLX (SC)) exhibited marginal issues with FDR control in larger samples. Overall, the findings from the bulk RNA-Seq simulations suggest that our ensemble strategy yielded notable improvements in model performance by enhancing true positive detection while effectively controlling false discoveries.

### 2.3 Ensemble models enhance interpretability of single-cell differential expression while maintaining power and FDR

Having demonstrated the superiority of ensemble-based differential expression in bulk RNA-Seq data, we next assessed whether the same approach is effective for single-cell RNA-Seq (scRNA-Seq) DE analysis. Similar to bulk RNA-Seq, we considered state-of-the-art models commonly used for the detection of DEGs in scRNA-Seq data (**Methods**) as the core. Three core models—DESeq2, edgeR, and VOOM—originated from the bulk RNA-Seq literature but are widely used in the single-cell community ^18,88^. Among other single-cell-specific methods, Tweedieverse operates under the compound Poisson GLM framework ^29^, and MAST uses a hurdle linear model ^24^. Following recent single-cell DE benchmarking efforts^29,88^, we also included metagenomeSeq ^75^, which employs a zero-inflated Gaussian linear model and was originally developed for microbiome data.

We used logistic regression (LR) as the enhancer in creating the two-part ensembles for three primary reasons (**Table 1**). First, in the context of single-cell differential expression, it has been consistently observed that a binarized representation of single-cell expression data more accurately captures biological variation and reveals the relative abundance of transcripts than raw counts^7,21^. Second, unlike differential gene expression analyses, which typically focus on testing differences in average gene expression between cell types or conditions, differential detection assesses differences in the fraction of samples or cells in which a gene is detected^35^. This is often evaluated using a logistic regression model based on binarized expression profiles^21^, an approach also prevalent in related fields such as ecology, microbiome research, and spatial omics ^41,64,70^, where presence–absence differences can be as meaningful as differences in mean values. Third, differential expression and detection analyses provide additive insights—both in terms of the specific genes they identify and their functional interpretation^35^. From a biological actionability standpoint, prioritizing genes that show differences in expression, detection, or both may be more informative, offering biologists a set of target genes with multimodal relevance. Notably, we did not consider ensemble versions of MAST and metagenomeSeq, as both methods already incorporate an LR component in their two-part formulations, rendering additional ensemble modifications redundant.

To evaluate these models across a range of scenarios, we generated synthetic data using the state-of-the-art scRNA-Seq data simulator, SCRIP^79^. SCRIP uses an existing scRNA-Seq dataset as a template to generate synthetic counts that capture key features of the original data, such as dropout rate, mean-dispersion trends, and technical and biological variability (**Methods**). Each simulated dataset contained 10, 000 genes, with 10% designated as DE genes. To assess model performance across both small and large sample sizes, we analyzed datasets containing a total of *N* = 200 and *N* = 500 cells, respectively, equally divided between two conditions. Additionally, we varied the differential expression factor to 1, 1.5, resulting in log2-fold changes that realistically represented both modest and strong effect sizes ^29,79,107^. We used a subset of a public human brain dataset (GSE67835^23^), available from the SC2P R package ^103^, as a template (**Methods**). Similar to our bulk RNA-Seq evaluations, we assessed: (i) the ability of the core models and their ensemble variants to detect DE genes (sensitivity/power), and (ii) their ability to control the FDR at the nominal 5% level (**Methods**), averaging results over 100 replications.

For cases with a smaller number of cells (*N* = 200) and a small effect size (differential expression factor = 1), as well as a larger number of cells (*N* = 500) and a large effect size (differential expression factor = 1.25), all core models and their ensemble variants successfully maintained FDR control at the 5% level (**Table 2**). All FDR-controlling ensemble methods demonstrated a significant boost in power compared to their non-ensemble cousins. Among the ensemble configurations, CC exhibited the smallest increase in power, followed by MC, while SC showed the highest increase. This trend was consistent across both small and large sample sizes and effect sizes. We note that while the SC method is not optimal for pooling dependent tests, it has been successfully applied in large-scale hypothesis testing for microarray data^42^, demonstrating favorable performance similar to our observations in this study. Despite these encouraging findings, further research is needed on the use of SC for pooling dependent hypotheses, as ignoring this dependence could lead to overly conservative or overly liberal Type I error rates ^15^. Overall, consistent with our findings from the bulk RNA-Seq evaluations, the scRNA-Seq results suggest that our ensemble strategy substantially improves performance, enhancing sensitivity while effectively controlling false discoveries in scRNA-Seq DE analysis.

**Table 2:**
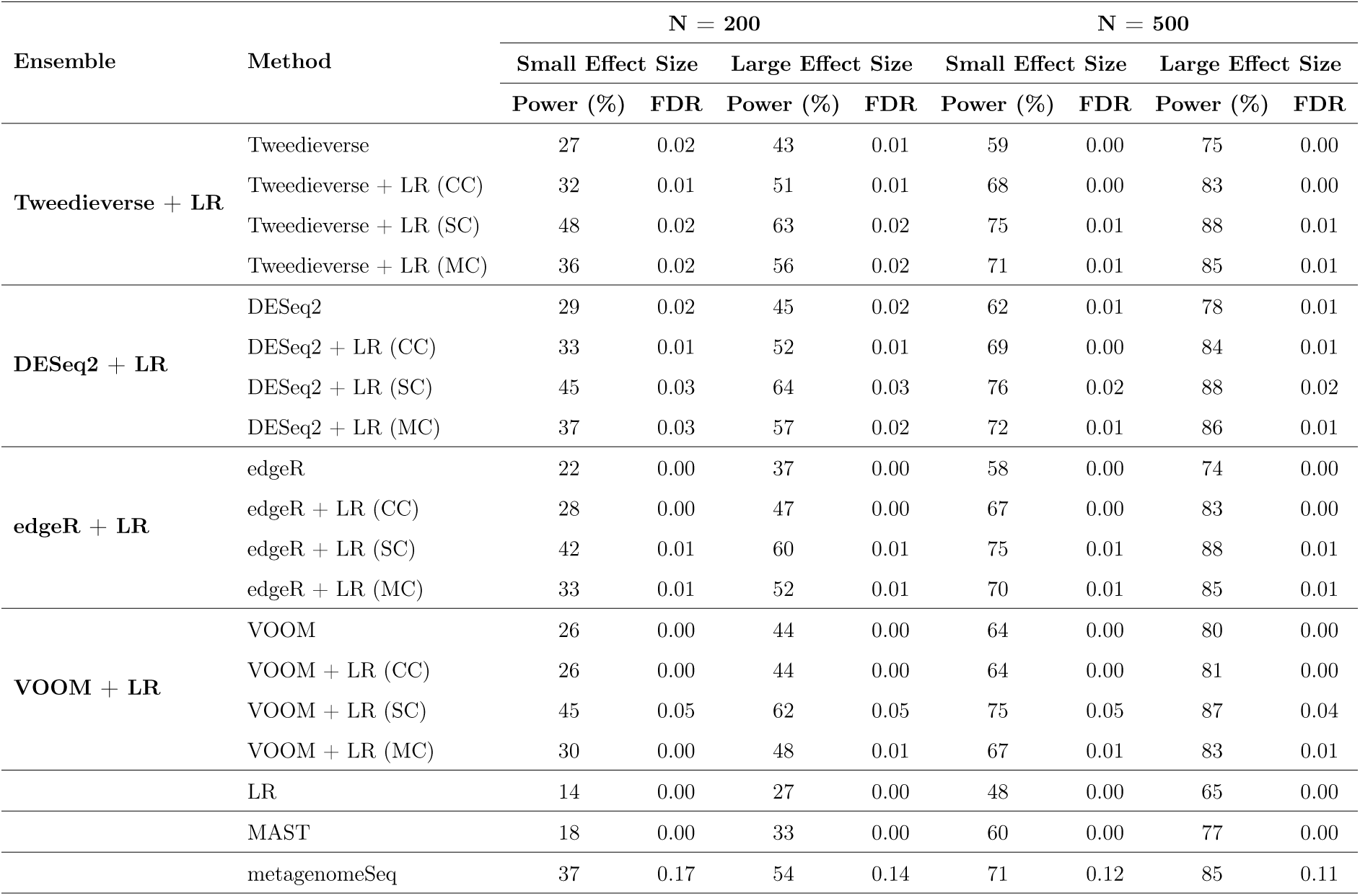
Power and FDR of ensemble models for scRNA-Seq DEG detection across varying simulation parameters. This table summarizes the power (true positive rate) and FDR for core models (Tweedieverse, DESeq2, edgeR, and VOOM), each combined with a logistic regression (LR) enhancer under three ensemble variants (CC, SC, MC), across two sample sizes (*N* = 200, *N* = 500) and two effect sizes (small and large). Standalone models (LR, MAST, and metagenomeSeq) are also included for comparison. All values represent averages over 100 replications.

### 2.4 Ensemble learning consistently enhances the precision and recall of microbiome differential abundance detection

Next, we examined the impact of ensemble learning on microbiome compositional data, focusing on differential abundance analysis, where controlling the false discovery is particularly challenging ^26,28^. Although conceptually similar to DE analysis, differential abundance analysis must account for the compositional and hierarchical structure of microbiome data, along with challenges common to traditional omics studies, such as high dimensionality, count measurements, sequencing depth variation, sparsity, and non-normality ^28^, among others. We applied the ensemble strategy to six core microbiome DAA methods (**Methods**; **Table 1**): ALDEx2^32^, ANCOM-BC2^52^, DESeq2^60^, LinDA ^113^, LOCOM ^27^, and MaAsLin2^28^. Most of these methods incorporate compositional adjustments through specialized normalization techniques or statistical corrections to address biases inherent in relative abundance data. LR was chosen as the enhancer for its simplicity, interpretability, and strong performance in prevalence modeling, which can potentially capture distinct biological signals from presence-absence patterns ^64,70,76^.

We evaluated the performance of microbiome-tailored ensemble models across four scenarios. For each scenario, we generated 100 synthetic microbiome datasets using SparseDOSSA ^62^, following recent benchmarking efforts^28,70^ (**Methods**). Specifically, we assessed (i) the ability of core models and their ensemble variants to detect differentially abundant features (recall/power) and (ii) their capacity to remain high-precision. Additionally, we included two standalone methods—metagenomeSeq ^75^ and MaAsLin3^70^—which inherently incorporate LR within their two-part formulations. Consistent with findings from non-microbiome simulations, microbiome ensemble models generally achieved higher recall than their standalone counterparts (**Figs. 3**; **S1**) while preserving precision (**Figs. S2–S3**). Notably, ALDEx2^32^ and MaAsLin2^28^ demonstrated substantial power improvements across all simulation scenarios, although no single ensemble variant consistently outperformed the others. These results reinforce the advantage of ensemble approaches in differential abundance analysis, highlighting their potential to enhance detection sensitivity while maintaining precision.

**Figure 3:**
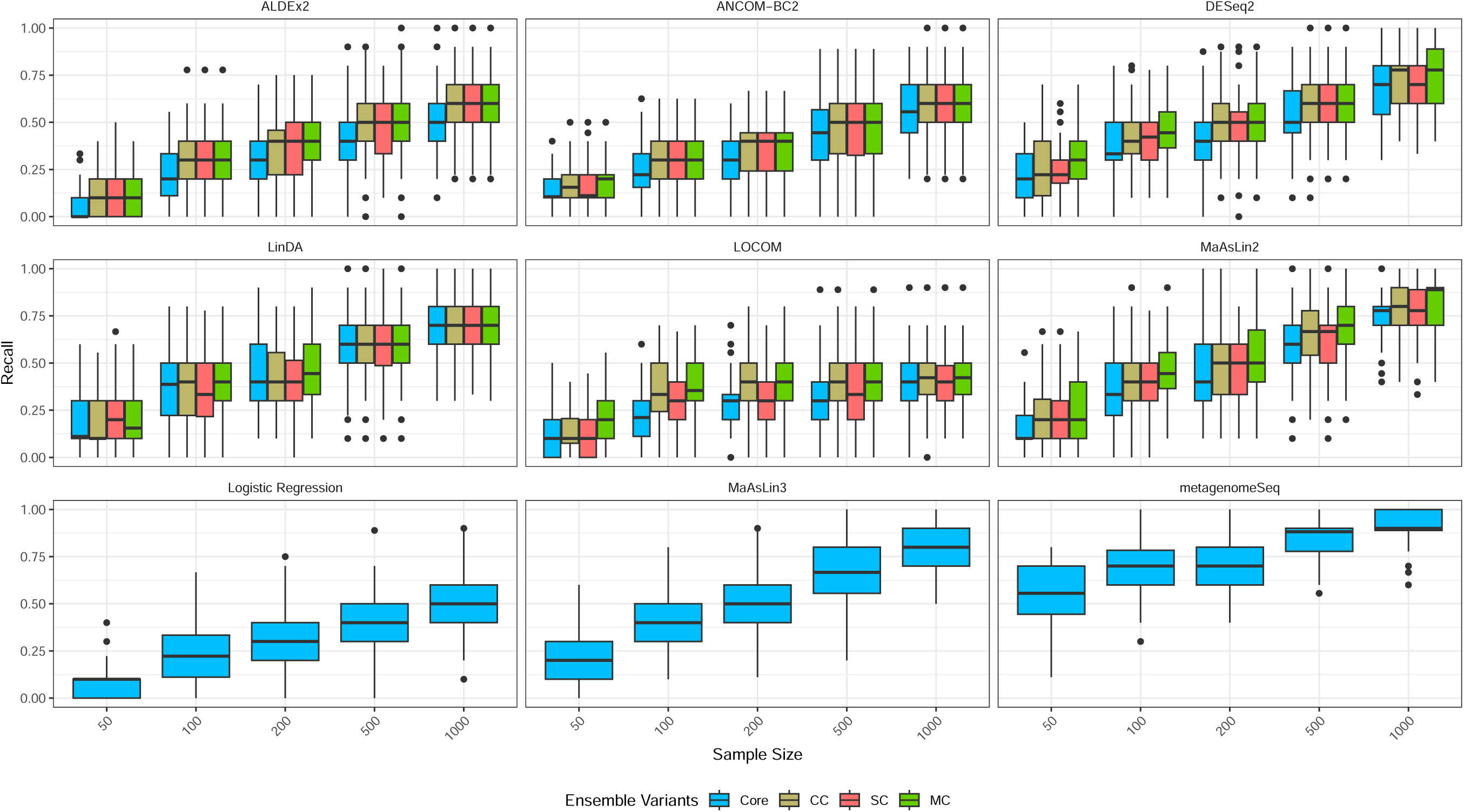
The impact of ensemble learning on differential abundance power across various simulation parameters for synthetic microbiome counts. We simulated 100 datasets with five sample sizes (50, 100, 200, 500, 1000) and effect sizes drawn uniformly from 2.5 to 5, with 10% of features being truly differentially abundant out of a total of 100 features and univariate binary metadata. Power is shown for six core microbiome analysis methods (ALDEx2, ANCOM-BC2, DESeq2, LinDA, LOCOM, MaAsLin2), each augmented with an LR enhancer using the CC, SC, and MC p-value combination strategies. Standalone models (LR, MaAsLin3, and metagenomeSeq) are also included for comparison. The ensemble approach consistently increased recall while maintaining precision across all simulation scenarios (**Figs. S1–S3**).

It is to be noted that several non-ensemble models proposed in the literature mirror the fundamental intuition behind ensemble models, with zero-inflated (metagenomeSeq) and hurdle classes of models (MaAsLin3, MAST) being notable examples ^24,70,75,105,109^. However, both classes of models suffer from critical interpretability issues as they do not directly model the feature-level population mean but instead rely on the conditional mean to test for differential association. Zero-inflated models, in particular, have regression coefficients with latent class interpretations, making them unsuitable for quantifying the effect of an explanatory variable on the feature-level population mean ^58,78^. Hurdle models, on the other hand, treat all zero observations separately from positive realizations, assuming that zeros arise from a distinct data-generating mechanism different from the process producing positive observations. Consequently, neither zero-inflated nor hurdle models offer a straightforward framework for interpreting and inferring covariate effects on the feature-level population—an interpretation that is almost always desirable in major biological applications. Although marginalized models have been proposed to address this limitation^11,58,78^, recent evaluations indicate that dedicated bespoke marginalized models provide no significant advantage over traditional non-marginalized models^53^. Moreover, marginalized models are often complex to solve algorithmically and unsuitable for large-scale applications, which reduces their practical utility ^53^. In contrast, our ensemble framework, by formulation, does not rely on a conditional modeling paradigm and instead provides directly interpretable parameter estimates, representing the overall covariate effect on the feature-level population.

### 2.5 Analysis of multi-domain omics datasets disclose biological discoveries that otherwise cannot be revealed by existing approaches

Following the proof-of-concept validation in synthetic benchmarking, we set out to examine whether the ensemble strategy could facilitate novel discoveries in published studies. To this end, we considered three key questions: (1) Can our ensemble strategy help resolve discrepancies between different non-ensemble DAA methods in cases where they tend to disagree? (2) Does it have the power to detect effects that some or most published methods might miss? (3) Can it uncover entirely new discoveries through data pooling by combining marginally significant effect sizes, similar to how boosting in machine learning combines weak learners to strengthen predictions ^9^? To answer these questions comprehensively, we reanalyzed seven published datasets across disease biology, spanning bulk, single-cell, and microbiome omics, demonstrating how the ensembling approach can effectively address the so-called ‘reproducibility crisis’ in published studies ^28,29,31,50,76^ and lead to both unique and consensus discoveries that are not achievable by standalone, non-ensemble approaches (full results available in the **Supplementary Materials**; **Supplementary Data S1-S7**).

#### 2.5.1 Analysis of Bulk RNA-Seq Data

To assess the performance of the two-part ensemble strategy on population-level bulk RNA-Seq data, we analyzed the liver cancer (LIHC) dataset ^50^, which contains 60,483 gene expression profiles from 100 matched healthy and cancer tissue samples. Five core models—edgeR, DESeq2, VOOM, Robseq, and dearseq—were used alongside their ensemble variants to identify DE genes from this dataset. Across all models, ensemble variants detected more DE genes compared to their non-ensemble counterparts (Fig. 4; **Supplementary Data S1**). Among the ensemble approaches, the SC variant consistently detected the highest number of DE genes, followed by the MC and CC variants, except for the dearseq family of methods (Fig. 4). For instance, edgeR + WLX (SC) identified 10,694 DE genes (compared to 8,595 for edgeR), 10,870 with DESeq2 + WLX (SC) (vs. 9,148 for DESeq2), 10,485 with VOOM + WLX (SC) (vs. 8,244 for VOOM), 12,846 with dearseq + WLX (SC) (vs. 11,684 for dearseq), and 10,808 with Robseq + WLX (SC) (vs. 8,975 for Robseq). These findings align with our simulation results, where all ensemble strategies demonstrated increased sensitivity in identifying DE genes compared to core models. Additionally, ensemble variants uniquely identified thousands of DE genes that were missed by the core models without the enhancer, with varying degrees of overlap across ensemble approaches (Fig. 4). As SC was the best performer in bulk RNA-Seq evaluations, we focus our discussion on the SC ensemble in the subsequent sections.

**Figure 4:**
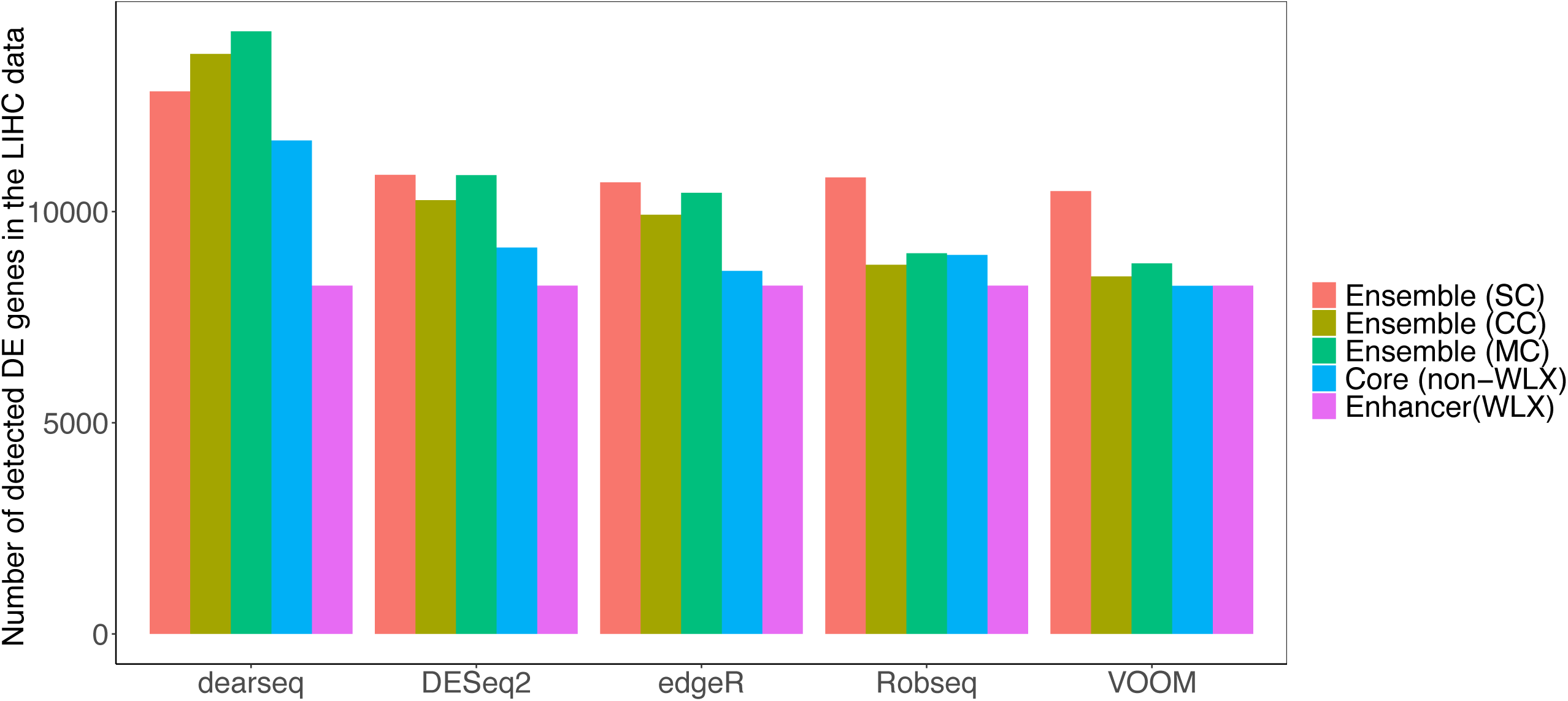
Comparison of differentially expressed genes detected by various DE models for the LIHC dataset^50^. We analyzed five core models (dearseq, DESeq2, edgeR, RobSeq, and VOOM) along with their ensemble counterparts, using the Wilcoxon (WLX) model as an enhancer. Ensembles incorporated three p-value aggregation methods—Cauchy Combination (CC), Stouffer’s Combination (SC), and minP Combination (MC)—to derive combined p-values followed by FDR adjustment. Genes with FDR < 0.05 were considered differentially expressed. Results show that ensemble models consistently identify more DE genes than individual models, highlighting their effectiveness in capturing complex biological signals.

Beyond the overall increase in DEG detection, two additional observations emerged. First, the ensemble strategies improved agreement between different DE models. The overlap of DE genes identified across all core models was 5,174, whereas the overlap across SC ensemble variants increased to 8,677, a 68% improvement (**Supplementary Data S8**). This suggests that ensemble methods help reconcile discrepancies among DEG detection tools, resulting in more consistent and robust findings. Second, ensemble variants identified DE genes with low effect sizes that were overlooked by the core models. For example, the SC ensemble variant of edgeR and DESeq2 identified 2344 and 1997 unique DE genes, with log fold changes ranging between (−1.1 and 1.2) and (−1.4, 1.3) respectively (**Fig. S4**; **Supplementary Data S9**). Similarly, Robseq’s ensemble variant detected 1,833 unique DE genes with effect sizes between −1.2 and 1.7 (**Fig. S4**; **Supplementary Data S9**). These findings indicate that ensemble strategies may enhance sensitivity, particularly for subtle expression changes.

To assess the biological relevance of the findings, we examined the top genes uniquely identified by *DAssemble*. We compiled DE genes that overlapped across different ensemble strategies (CC, MC, SC) and selected the top 10 based on adjusted p-values (**Supplementary Data S10**). Among them, SERPIND1, a gene encoding heparin cofactor II, was identified. SERPIND1 is a serine protease inhibitor involved in blood coagulation and has been reported to show altered expression in hepatocellular carcinoma (HCC), potentially influencing tumor progression and metastasis^99^. Another notable gene, GABPA, encodes a subunit of the GA-binding protein transcription factor, which plays a role in various cellular processes, including cell cycle progression and mitochondrial function ^108^. In HCC, GABPA expression is significantly reduced in tumor tissues compared to adjacent normal tissues, correlating with elevated alpha-fetoprotein levels, advanced tumor grade, and distant metastasis^108^. Functionally, GABPA inhibits HCC cell migration and invasion both in vitro and in vivo, partly by regulating E-cadherin, a protein essential for cell-cell adhesion whose loss is associated with increased tumor invasiveness^39^. By modulating E-cadherin expression, GABPA functions as a tumor suppressor, limiting the metastatic potential of HCC cells ^39,108^. Notably, these biologically relevant signals were missed by all core models but were uniquely captured by *DAssemble*, reinforcing the potential of ensemble strategies to detect meaningful molecular signals in RNA-Seq analyses.

As a final RNA-Seq application, we analyzed a GTEx dataset ^59^, consisting of 56,200 genes and 758 cells collected from heart tissue (**Table S1**). Similar to previous analyses, all three ensemble approaches detected more DE genes than their respective core models in this dataset (**Fig. S5**; **Supplementary Data S2**). Additionally, the ensemble approach identified many unique DE genes, expanding the hypothesis space for potential biological discoveries beyond what individual models could detect. These findings further support the robustness of ensemble strategies in improving DE gene detection across varied RNA-Seq datasets, both small and large.

#### 2.5.2 Analysis of Single-cell RNA-Seq Data

Having demonstrated the clear advantage of ensemble learning in bulk RNA-Seq DE analysis, we next examined whether similar performance gains could be observed in single-cell differential expression analysis. To this end, we analyzed heterogeneous scRNA-seq data from two broad categories of scRNA-seq protocols ^29^: unique molecular identifiers (UMIs) and read counts (non-UMI). For our primary demonstration, we focused on a non-UMI dataset referred to as the Brain data, which includes expression counts for 100 single cells (*N* = 38 oligodendrocytes and *N* = 62 astrocytes) across 10,483 genes ^23,102^. We further analyzed 1) one UMI dataset containing 13,713 genes and 2,638 peripheral blood mononuclear cells (PBMCs)^111^, and two additional non-UMI datasets from mouse kidney cells ^73^ and human preimplantation embryonic cells^77^ (**Table S1**; **Supplementary Data S3-S6**).

In contrast to bulk RNA-seq, scRNA-seq has been shown to have a higher fraction of zeros per sample, which can be a primary source of variation and influence downstream analyses ^29^. We thus extended the previous analysis of bulk RNA-seq gene expression by focusing on single-cell-specific DE methods that have consistently performed well in recent benchmarks^29^, using four core models (edgeR, DESeq2, VOOM, and Tweedieverse), along with their LR-enhanced two-part ensembles (**Table 1**). Similar to before, we also considered a standalone method (MAST ^24^) that already includes an LR component and was used without ensemble modification.

As before, we focus on the SC ensemble due to its strong performance in simulations (**Table 2**), highlighting its enhanced sensitivity and robustness in detecting differentially expressed genes. As expected, ensemble models identified a substantially higher number of DE genes compared to the non-ensemble models (Fig. 5A). For instance, edgeR detected 1,118 DE genes, whereas its single-cell ensemble variant identified 1,755. A similar trend was observed for DESeq2 (1,553 vs. 2,105), VOOM (1,618 vs. 2,440), and Tweedieverse (1,331 vs. 2,034). In contrast, MAST identified 1,350 DE genes, fewer than the two-part ensembles. This phenomenon was consistent across all the datasets we analyzed (**Figs. S6-S11**), underscoring the advantage of *DAssemble* in improving sensitivity while maintaining robust differential expression detection across diversified single-cell datasets.

**Figure 5:**
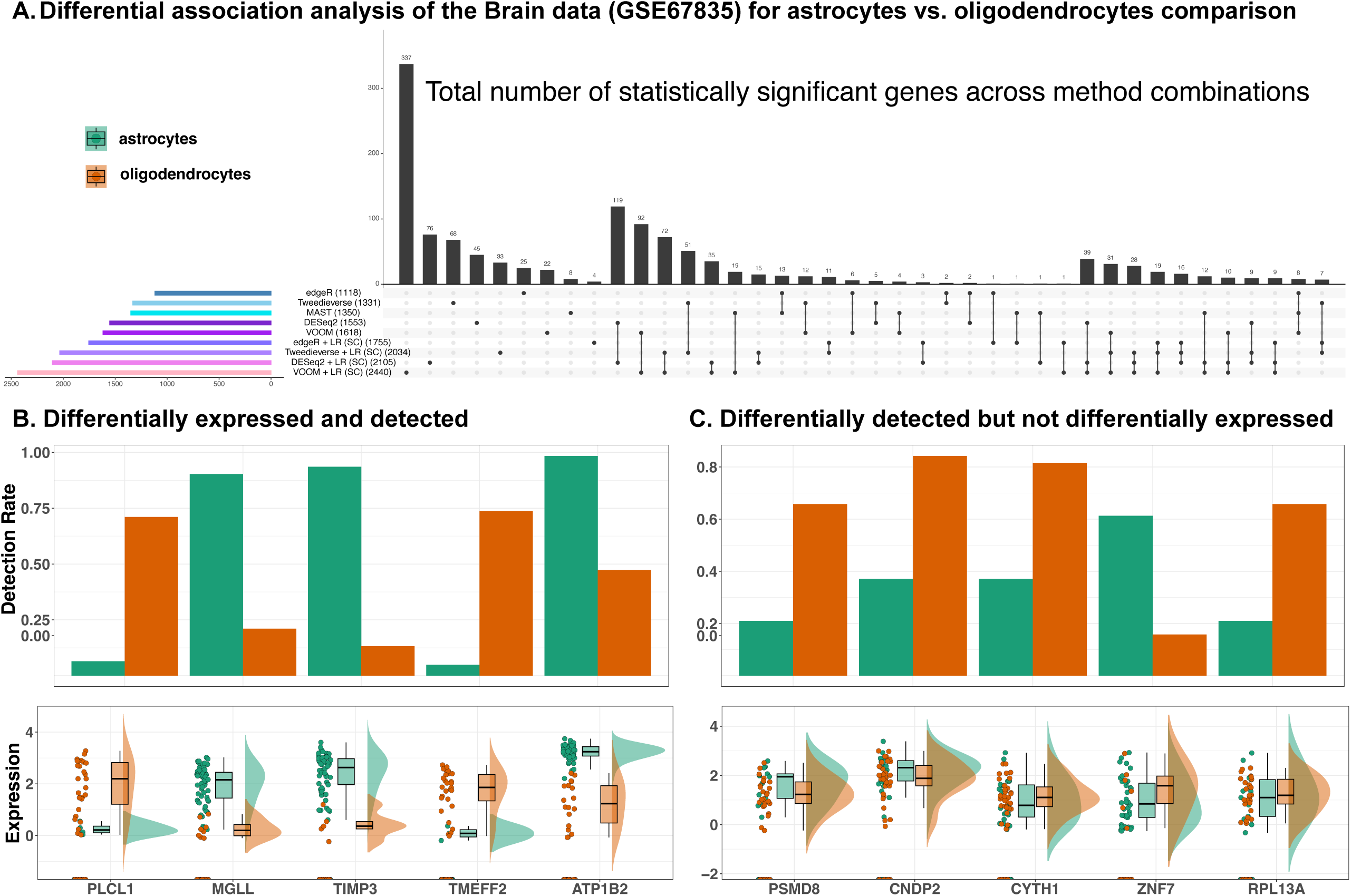
Differential analysis of scRNA-Seq profiles comparing astrocytes and oligodendrocytes reveals new biological insights enabled by the ensemble approach. ^23^**. A.** UpSet plot showing the overlap in differentially expressed genes (DEGs) identified across methods, comparing astrocytes (*N* = 62) and oligodendrocytes (*N* = 38). Horizontal bars indicate the total number of DEGs per method; vertical bars show shared DEGs across method combinations. **B.** Raincloud plots for the top five DEGs that were statistically significant in both differential expression and detection. **C.** Raincloud plots for the top five DEGs that were significant in detection only. Detection rates are shown above, with expression distributions across celltypes shown below. The Tweedieverse + LR (SC) ensemble was used to select representative genes.

Crucially, the impact of ensemble learning is evident not only in the number of DEGs detected but also in their biological significance. In particular, we identified two sets of previously unknown DEGs using the Tweedieverse + LR (SC) ensemble as a representative approach. The first set, differentially expressed and detected genes, reveals potentially novel biological mechanisms, as their expression differs between astrocytes and oligodendrocytes not only in terms of average expression but also in detection rate (Fig. 5B). These genes may serve as key regulatory factors, such as transcription factors or signaling molecules, shaping cell identity by influencing both expression magnitude and the proportion of cells in which they are active ^85,110^. Their simultaneous differential expression and detection suggest an integrated mode of regulation, possibly driven by lineage-specific transcriptional programs or epigenetic modifications^61^. Furthermore, these genes could play a role in fundamental astrocyte-oligodendrocyte interactions, including metabolic support, myelination processes, and neuroinflammatory responses ^34,40^. Their distinct presence across both expression and detection metrics underscores their potential as master regulators of glial cell function^93^.

The second set of genes, differentially detected but not differentially expressed, are identified as significant by the ensemble model for their highly significant differential detection rate across celltypes (Fig. 5C). Notably, they remain undetected by the core DE methods, showing no variation in average expression between celltypes (Fig. 5C). This unique category of genes may reflect transcriptional heterogeneity within a given cell type, where expression is restricted to specific subpopulations rather than being ubiquitously present^43,83^. Such genes could play roles in specialized cellular functions, responding to external stimuli or intrinsic regulatory cues that modulate their activation in a subset of cells ^92^. Alternatively, their differential detection may result from stochastic transcriptional bursts, where gene expression is temporally regulated rather than continuously maintained, leading to fluctuations in the proportion of cells expressing these genes at any given time ^45,80^. Another possibility is that these genes are influenced by threshold-dependent regulatory mechanisms, wherein subtle differences in transcription factor binding or chromatin accessibility push certain genes above or below the detection threshold in a subset of cells ^8,54^.

It is clear that an ensemble approach that integrates multiple biologically informative signals has the potential to uncover novel regulatory mechanisms that traditional differential expression analyses—reliant on strict fold change or mean expression thresholds—may overlook. While our two-part formulation employed logistic regression for differential detection, incorporating additional biologically motivated strategies (e.g., a three-part ensemble to simultaneously assess differential variability ^38^, expression, and detection) could further enhance discovery. Taken together, our ensemble framework provides a general approach for uncovering hidden layers of gene regulation by expanding the hypothesis space and increasing the likelihood of discovery, moving beyond mean expression changes toward a comprehensive, multimodal understanding of gene expression across varied biological contexts.

#### 2.5.3 Analysis of Microbiome Data

The usefulness of our ensemble approach, of course, is not limited to gene expression studies. Moving beyond univariable differential expression analysis, we now apply our ensemble strategy to the multivariable differential abundance analysis of a population-level microbiome multi-omics study, where the challenges are amplified. Unlike gene expression studies, where read counts can be analyzed more straightforwardly without severe compositional constraints, microbiome multi-omics requires careful handling to account for relative abundances, sparsity, and confounding factors such as host demographics, clinical variables, and dietary influences^28,65^.

To fully assess the impact of ensemble learning on microbiome differential abundance detection, we systematically evaluated microbial associations in the integrative Human Microbiome Project (iHMP) ^57^. Specifically, we leveraged the baseline multi-omics data from the iHMP Inflammatory Bowel Disease Multi-omics Database (IBDMDB). IBDMDB includes 132 individuals recruited from five U.S. medical centers^57^ followed longitudinally for one year with up to 24 time points per individual. The baseline cohort comprises participants diagnosed with Crohn’s disease (CD, N = 41) and Ulcerative Colitis (UC, N = 30), jointly grouped as IBD (N = 71), as well as non-IBD controls (N = 23). The multivariable model included IBD disease phenotype (with non-IBD as the reference), age, and antibiotic use as fixed effects. This allowed us to test the validity of the ensemble approach in more complex modeling scenarios, particularly in the presence of multiple covariates and compositionality.

Following the preprocessing and quality control procedures outlined in previous studies^28,70^, we applied seven core microbiome methods (MaAsLin2, LinDA, ANCOM-BC2, ALDEx2, LOCOM, edgeR, and DESeq2) to the iHMP microbial taxonomic profiles (**Methods**) based on MetaPhlAn 4^6^. For models requiring count data (LOCOM, DESeq2, edgeR, and ANCOM-BC2), we first transformed relative abundances into approximate read counts before model fitting^70^. To apply the ensemble strategy, we extracted the IBD-specific associations and p-values from the corresponding feature-metadatum pairs in the multivariable model, then applied three LR-enhanced ensemble strategies (CC, MC, and SC) as before, followed by FDR correction (**Figs. 6; S12-S14**).

**Figure 6:**
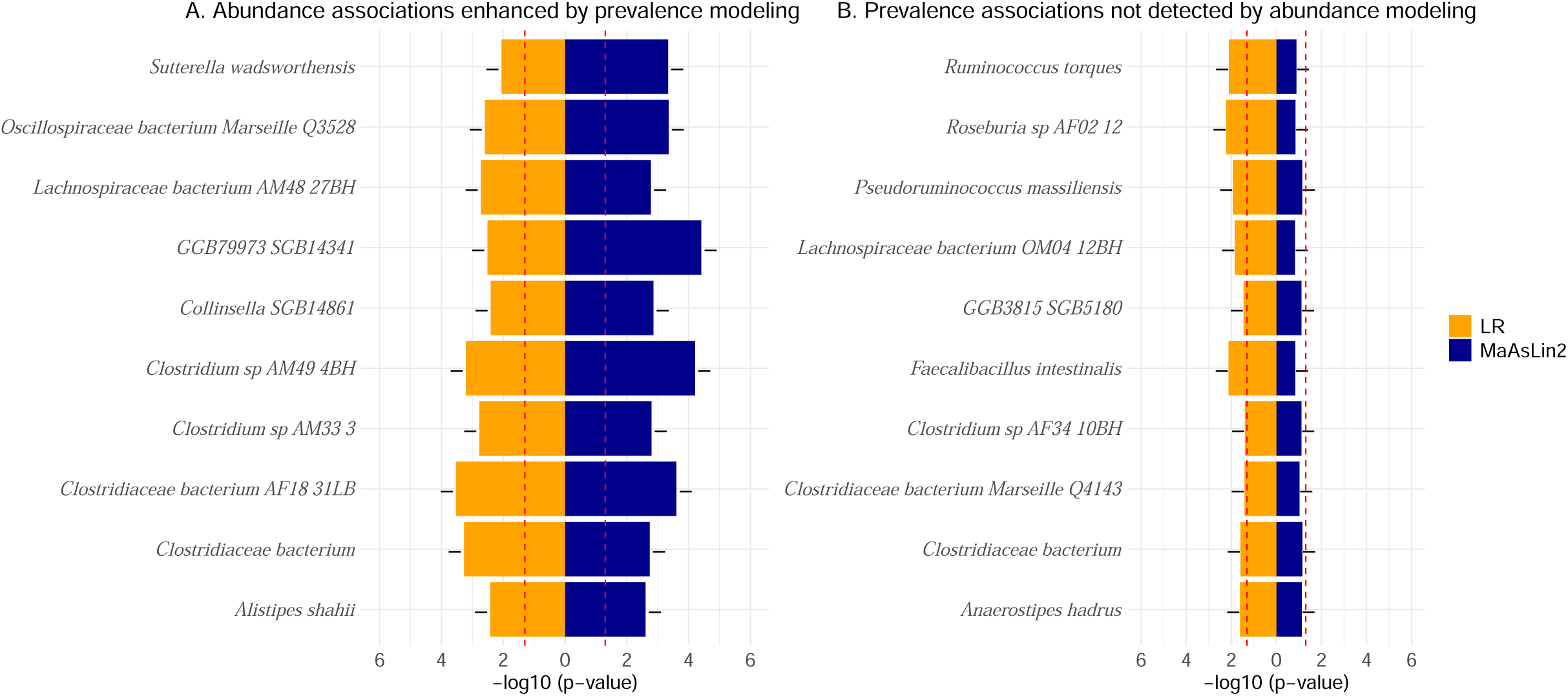
Ensemble modeling reveals additional microbial species associated with IBD status beyond conventional methods. We used MaAsLin2 and LR as the base and enhancer models, respectively, along with the SC ensemble approach as a representative method for the iHMP data. Here, we report the top 10 microbial species ranked by adjusted p-values from the SC ensemble, which integrates p-values from MaAsLin2 and LR before FDR correction. The left panel compares log_10_(*p*) values from MaAsLin2 and LR for microbial species identified as significant by both methods. The right panel highlights species detected only by the ensemble despite being ranked poorly by MaAsLin2—demonstrating the enhanced sensitivity of the combined approach. Plus (+) and minus (–) signs indicate the direction of association with IBD. These results underscore the benefit of ensemble learning in uncovering microbial associations that may be overlooked by individual models. A red line in these plots represents the nominal type-I error rate, set at 0.05.

There is a clear statistical and biological rationale for selecting LR as the enhancer. Biologically, the presence or absence of microbiome features can distinguish distinct processes, such as infection versus commensal activity ^64,70^. Statistically, prevalence modeling mitigates biases from pseudo-count transformations that affect abundance modeling in common DAA methods ^70^. Unlike hurdle models ^70^, which model only nonzero abundances in the abundance component, and zero-inflated models ^75^, which rely on untestable latent-class assumptions ^58,78^, our two-part ensemble approach accounts for both overall abundance and prevalence, leading to marginal inference without conditioning ^58,78^. By preserving zeros in the abundance model, this approach better captures the true distribution of microbial features across samples, offering a more interpretable framework for inferring the population mean, potentially enhancing the explainability of disease-associated microbiome shifts across the population.

Several noteworthy observations emerged. The recurring theme of an increase in discovery by *DAssemble* was recapitulated as before, though to a lesser extent in microbiome data analysis, as some ensemble approaches did not enhance discovery (**Fig. S14**). This is potentially due to compositional effects canceling out abundance- and prevalence-driven associations ^70^, suggesting that alternative compositionally aware aggregation methods may be beneficial for microbiome-based ensemble approaches. Notably, the original publication reported “no statistically significant associations at baseline,” albeit using a different version of the software (MetaPhlAn 2^94^) and a slightly different statistical model (MaAsLin2^28^). In synergy, both LOCOM and LinDA did not yield statistically significant cross-sectional associations, reinforcing the idea that the two examined groups (IBD and non-IBD) are strikingly similar ^57^, making the analysis challenging with potentially many small effects (**Supplementary Data S7; Fig. S12**).

Despite the challenges, the ensemble strategy improved sensitivity in detecting biologically relevant microbial associations. Using MaAsLin2 + LR (SC) as a representative method, we identified two major classes of IBD-associated taxa: 1) exhibiting both broad compositional shifts and presence-absence patterns (Fig. 6A); and 2) prevalence associations not detected by abundance modeling alone (Fig. 6B). Notable species where abundance-based associations were enhanced by prevalence modeling include *Roseburia* species and *Pseudoruminococcus massiliensis*. *Roseburia* species are recognized for their butyrate production and anti-inflammatory properties, and their depletion in IBD patients has been well-documented, suggesting a protective role in gut homeostasis ^63^. Similarly, *Pseudoruminococcus massiliensis* has been identified as a butyrate-producing bacterium that plays a role in gut health and metabolism, though its direct involvement in IBD is still being explored^37^. *Collinsella* species have been reported to modulate gut inflammation, with higher abundance linked to altered metabolic pathways in IBD patients^71^. Likewise, *Alistipes shahii* has shown variable associations with gut inflammation, with some studies suggesting a potential link to disease exacerbation^74^.

Conversely, notable species whose prevalence-based associations were completely missed by abundance modeling include *Anaerostipes hadrus* and *Faecalibacterium intestinalis*, both of which have been underrepresented in IBD patients. *Anaerostipes hadrus*, another butyrate producer, has shown decreased prevalence in individuals with IBD, reinforcing its protective role against intestinal inflammation^112^. While specific studies on *Faecalibacterium intestinalis* are limited, the closely related *Faecalibacterium prausnitzii* is one of the most well-characterized anti-inflammatory bacteria in the gut microbiome. *F. prausnitzii* plays a critical role in maintaining gut health through butyrate production and immune modulation, and its depletion has been consistently observed in IBD patients^57,112^. Given their phylogenetic proximity, *F. intestinalis* may share similar protective properties, though further research is needed to clarify its specific role in IBD. These findings underscore the importance of integrating both abundance and prevalence modeling in microbiome research to fully capture microbial contributions ^70^.

Beyond abundance- and prevalence-based associations, *DAssemble* can potentially be extended to conduct a differential distribution analysis ^14^ of microbiome data by incorporating other reinforcing characteristics, such as dispersion^14^ or dysbiosis ^68^ - an advancement not straightforwardly achievable with existing methods. By capturing multiple dimensions of microbial variation, this approach can detect subtle yet biologically meaningful shifts in microbial communities. This is particularly critical in the context of complex human diseases, where heterogeneous microbial alterations and low signal-to-noise ratios may drive disease progression—underscoring the need for integrative methods to fully unravel microbiome-health associations.

## 3 Discussion

Ensemble learning is a popular and powerful tool in machine learning. In this paper, we develop a framework that repurposes the power of ensemble learning for a simpler statistical setting: differential association analysis. By integrating diverse base models, our ensemble approach circumvents the limitations of selecting a single optimal model and leverages the strengths of complementary statistical approaches, thereby enhancing the detection of differential patterns while ensuring reliable control of false positive and false discovery rates. Beyond DAA, the ensemble strategy is agnostic to the statistical task at hand and can be extended to various downstream analyses, including normalization, differential network analysis, mediation analysis, batch effect correction, gene set enrichment analysis, and meta-analysis, among others. Our findings thus have broad implications for the field, particularly in harmonizing disparate statistical methodologies and fostering greater consensus in omics data science ^26,28,29,31,50^.

We note that our proposed framework can be considered a generalization of the ensemble strategy described in Liu *et al.* ^55^, which was developed for WGS association studies for testing a global null. Except for hurdle models ^24,70^, all existing multi-model frameworks, including scDEA^49^, fall into this category. Unlike Liu *et al.* ^55^, which focuses on testing a global null, our approach constructs a composite null by stacking multiple statistically and biologically enhancing null hypotheses, ensuring both statistical rigor and biological interpretability. In doing so, our framework clarifies the conceptual links among various versions of ensemble models by generalizing published two-part and hurdle methods and unifying them within a broader ensemble-based framework for differential association analysis. While our empirical results support its effectiveness, a formal theoretical justification for the composite null framework remains an important direction for future research.

While the ensemble approach offers numerous advantages, it also introduces certain complexities. For instance, computational costs can be a consideration for researchers working with large datasets or limited resources. We discuss many multi-type applications of this paradigm, including differential expression analysis of bulk and single-cell transcriptomics and differential abundance analysis of microbiome metagenomics. There are likely to be other interesting applications of these ideas, including epigenomics, proteomics, metabolomics, and lipidomics, as well as their integration in multi-omics studies^30^. While our study highlights the benefits of combining biologically motivated models with compatible characteristics, ensemble selection should be tailored to the dataset and the biological questions at hand^10^.

Consequently, this work presents several directions for future research. First, while we have focused on a fixed number of small candidate models in this manuscript, the general idea can extend to scenarios with many models, albeit at the expense of interpretability. Second, as omics studies advance toward higher spatial and temporal resolutions, we expect ensemble models to become even more critical for capturing complex spatiotemporal patterns. For example, in spatial transcriptomics, there is no clear consensus on the definition of a spatially variable gene (SVG), which refers to genes that exhibit non-random spatial patterns, and ensemble models might help mitigate the uncertainty associated with SVG identification^86^. Third, devising an approach similar to targeted learning ^18,95,97^—which combines flexible ensemble learning with a targeting step to refine initial estimates and align them closely with the target parameter—could be an important step in solidifying the statistical validity and interpretability of our framework. Fourth, selecting dynamic feature-varying candidate models, rather than relying on a single static model for all features, could better accommodate variability in feature-wise distributions ^16,29^. Automating the ensemble framework through self-supervised learning and incorporating it within multimodal AI agents^1,104,114^ would further enable multi-strategy biomedical applications, particularly for emerging omics modalities such as spatial multi-omics and digital pathology. Finally, exploring alternative p-value aggregation methods ^44,70,81,101^ may further validate the utility and generalizability of this approach across the broader omics research landscape.

In summary, the ensemble learning framework represents a significant advancement in differential analysis, offering a powerful and flexible tool for improving both current and future differential analysis methods while maintaining rigorous control over false positives. We envision this framework becoming a cornerstone in developing more robust and reproducible omics analyses, ultimately leading to more reliable biological insights and discoveries, much as has been the case for ensemble methods in machine learning ^30,96^. By significantly outperforming published methods across multifaceted datasets—including single-cell and bulk RNA-Seq as well as microbiome profiles—the ensemble approach provides a versatile, platform-agnostic solution for well-powered differential analysis of complex, high-dimensional omics data.

## Author Contributions

HM conceived the original idea, designed the numerical experiments, and provided overall supervision for the project. SC and EP conducted the bulk and single-cell analyses using both synthetic and real data. SC, EP, PB, and AB contributed to the interpretation of the results. ZL and JG performed the microbiome analyses using both simulated and real data and developed the associated software, with tutorial contributions from EP and CB. EP and HM coordinated and supervised the writing of the manuscript. All authors contributed to writing the final manuscript.

## Code Availability

The implementation of *DAssemble* is publicly available with source code, documentation, tutorial, and as an R package at https://github.com/himelmallick/DAssemble. Analysis scripts for the simulations (as described in Sections 2.2, 2.3, 2.4) and the real data analyses (as described in Section 2.5) are available from the corresponding author upon request.

## Acknowledgments

The authors gratefully acknowledge the resources provided by the high-performance computing (HPC) support and CPU time on the Cayuga compute cluster, made available by Information Technologies & Services (ITS) at Weill Cornell Medicine (WCM). We also extend our gratitude to Lilly Endowment, Inc. for its support of the Indiana University Pervasive Technology Institute. Additionally, we acknowledge Dr. Jia (John) Kang (Merck & Co., Inc., Rahway, NJ, USA) for providing initial feedback on an earlier version of the method.

## Methods

### Differential Analysis Methods

We applied several state-of-the-art DAA methods to both synthetic and real datasets. Unless otherwise noted, all tools were executed using their default settings. To ensure consistency across methods, we did not rely on the FDR-adjusted p-values provided by individual tools. Instead, we extracted the raw (unadjusted) per-feature p-values and applied a standardized FDR correction post hoc. Any deviations from default configurations are detailed below.

- **ALDEx2**^32^: We used the aldex() function from the ALDEx2 R package, which generates Monte Carlo samples from Dirichlet distributions with a uniform prior for each sample, performs centered log-ratio (CLR) transformations, and applies t-tests on the transformed realizations.
- **ANCOM-BC2**^52^: The ancombc2() function from the ANCOMBC R package was used with settings (*prv_cut* = 0, *struc_zero* = T), with other parameters left at their defaults. ANCOM-BC2 encountered substantial computational challenges; features that triggered errors during model fitting were excluded, and the model was subsequently refit.
- **dearseq** ^25^: Filtering and normalization followed the edgeR preprocessing pipeline described below. We then used the dear_seq() function from the dearseq R package with the asymptotic test for detecting differentially expressed (DE) genes.
- **DESeq2**^60^: Functions from the R package DESeq2 were used. First, a DESeqDataSet object was created from the count matrix using DESeqDataSetFromMatrix. Next, geometric means were manually computed from the raw counts and supplied to estimateSizeFactors(), followed by calling DESeq() for differential testing.
- **edgeR**^82^: Functions from the R package edgeR were used. We followed the standard edgeR pipeline. A DGEList object was created from the raw count matrix, normalization factors were computed using calcNormFactors() with the default Trimmed Mean of M-values (TMM) method, dispersions were estimated via estimateDisp(), and differential testing was performed using the likelihood ratio test via glmLRT().
- **VOOM** ^46^: Functions from the R package limma ^87^ were used. Feature counts were transformed using VOOM and analyzed using lmFit().
- **LinDA**^113^: We used the linda() function from the LinDA R package with default settings. LinDA is a linear modeling framework specifically designed for microbiome compositional data. It performs feature-wise linear regression on CLR-transformed data, followed by a compositional bias correction.
- **Logistic regression** ^17^: We implemented per-feature logistic regression using the glm() function from base R with a binomial family and logit link. Prior to modeling, data were binarized to indicate presence/detection (non-zero) or absence/non-detection (zero) of each feature.
- **LOCOM**^27^: We applied the LOCOM() function from the LOCOM R package using its default configuration. LOCOM is a logistic regression-based method specifically designed for compositional microbiome data. It models the log-ratio of each taxon’s relative abundance against a reference taxon and tests for differential abundance by comparing logistic regression coefficients between groups.
- **MaAsLin2**^28^: We used the MaAsLin2 R package with default settings. The method performs total sum scaling (TSS), replaces zeros with half the minimum observed value, applies a log transformation, and fits linear models on the transformed values.
- **MaAsLin3**^70^: We used the MaAsLin3 R package with default settings and median correction enabled. The pipeline includes (1) TSS normalization, (2) prevalence profiling while retaining non-zero abundances, (3) base-2 log transformation, (4) augmented logistic regression on the prevalence profile and linear regression on the non-zero abundance profile, and (5) effect size combination for each feature–metadatum pair.
- **metagenomeSeq** ^75^: We used the metagenomeSeq R package to create an MRexperiment object via newMRexperiment(), performed cumulative sum scaling (CSS) normalization using cumNorm() with p = 0.5, and conducted differential analysis using fitFeatureModel().
- **RobSeq** ^22^: We applied the RobSeq R package with default settings. This method uses edgeR-style preprocessing (TMM normalization and CPM filtering) followed by a robust linear model with degrees-of-freedom adjustments to the Welch t-statistic, following Bell-McCaffrey’s approach.
- **Tweedieverse** ^29^: We used the Tweedieverse R package with default settings. The method first applies an appropriate normalization procedure, then fits a per-feature Tweedie compound Poisson GLM. The Tweedie index parameter, which governs the variance function, is estimated individually for each feature from the data.
- **Wilcoxon rank-sum test**^67^: We applied the nonparametric Wilcoxon rank-sum test to each feature using the wilcox.test() function in base R. Prior to testing, we followed a standard preprocessing pipeline, including library size normalization. The test was then performed on the normalized values to assess group-wise differences.

### Simulation Details

To improve the robustness of our results and reduce random sampling errors, each simulation scenario was repeated 100 times. Features with low prevalence (<10%) were removed from the generated datasets to ensure robustness in the analysis.

### Bulk RNA-Seq Simulation

We used SimSeq ^2^ to generate synthetic data for our bulk RNA-Seq simulation studies. SimSeq is a nonparametric RNA-Seq data simulator that creates synthetic RNA-Seq read count matrices by sub-sampling from an existing template dataset. The simulation starts with a representative dataset, normalized using the TMM method from the edgeR^82^ R package. Each gene must be expressed in at least one sample from the groups under study. DEGs are selected via probability sampling, with preference given to genes with lower FDRs, based on p-values from the Wilcoxon rank-sum test. For equivalently expressed genes, samples are chosen using equal probability sampling from genes not identified as DE. The number of samples selected is balanced to ensure fair representation of both groups, enabling a thorough gene analysis.

### Single-cell RNA-Seq Simulation

We used SCRIP^79^ to generate synthetic data for our scRNA-Seq simulation studies. SCRIP is a flexible parametric simulator for scRNA-Seq data, designed to generate realistic synthetic datasets for various biological conditions. It captures key features of single-cell data, including gene expression variability, transcriptional bursting, and technical noise, and allows users to simulate data for multiple celltypes with control over parameters such as the number of cells, genes, and sequencing depth. Furthermore, SCRIP supports the simulation of differential expression between cell populations, enabling the exploration of biological hypotheses and facilitating robust benchmarking of computational methods for scRNA-Seq data analysis by incorporating both biological and technical variability.

### Microbiome Simulation

We used the R package SparseDOSSA2^62^ to generate synthetic microbiome count data. SparseDOSSA2 models marginal microbial feature abundances using a zero-inflated lognormal distribution and incorporates additional components to account for absolute cell counts, sequencing read generation, and both microbe–microbe and microbe–environment interactions. To generate realistic microbiome data, we used the “Stool” template provided by SparseDOSSA2, which captures key characteristics of real stool microbiome profiles, with read depths set to 50000. Unlike the non-microbiome simulators, SparseDOSSA2 enables the explicit injection of true positive associations at both the abundance and prevalence levels, modeled using a log-linear and a logistic framework, respectively.

### P-value Aggregation

In order to leverage the strengths of multiple statistical models, we aggregated p-values across methods to obtain a single omnibus p-value per feature. Specifically, assuming *N* observations, *P* features, and *K* candidate models *M*_1_*, M*_2_*, …, M_K_*, for any given feature *j*, we obtain a *K*-dimensional vector of p-values **p***_j_* = *p_jk_*, where each *p_jk_* corresponds to the p-value for feature *j* from model *k*. By applying appropriate p-value combination techniques, we summarize this vector into a single aggregated p-value *p*^0^ that reflects consensus evidence across models.

**Cauchy Combination Test (CCT):** The CCT provides an analytic approximation to the aggregated p-value using the Cauchy distribution, which is highly accurate even under arbitrary dependency structures among the input p-values ^56^. For the *j*^th^ feature, the combined p-value is computed as:

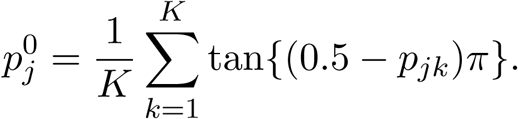

While CCT is appealing due to its simplicity and generality, it is sensitive to extremely large p-values: as any *p_jk_* → 1, the aggregated *p*^0^ tends to 1. This can be problematic in scenarios involving discrete data or model misspecification. To mitigate this, we employ a truncated version: where *ɛ* = 0.01 ensures numerical stability by capping the contribution of extreme p-values.

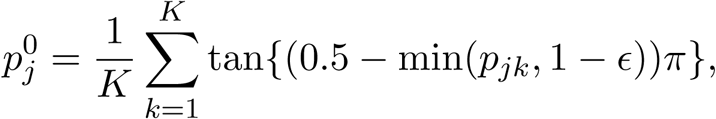
**Minimum P-value (minP):** The minP method ^12^ offers a simple alternative by selecting the smallest p-value across all *K* models:

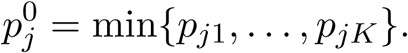

This approach emphasizes the strongest signal among the models, though it may be overly optimistic if not adjusted for multiple comparisons ^36^.
**Stouffer’s Method:** Stouffer’s method combines p-values by transforming them to the standard normal scale and averaging the resulting z-scores^90^. For the *K* p-values *p*_1_*, p*_2_*, …, p_K_*, the test statistic is:

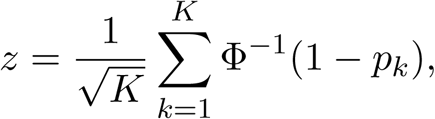

where Φ*^−^*^1^(.) is the inverse cumulative distribution function of the standard normal distribution. The aggregated p-value is then obtained from the resulting standard normal distribution.

Taken together, these p-value aggregation strategies allow us to synthesize evidence from multiple modeling approaches, enabling more robust and interpretable assessments of feature-level significance across heterogeneous methods.

### Performance Metrics

To evaluate the effectiveness of different methods, we considered two widely used performance metrics derived from the confusion matrix: Power and FDR (defined below). We assessed both the maximum and average values of these metrics across simulation replicates. In particular, power reflects the method’s ability to correctly identify truly significant features, while FDR quantifies the proportion of false discoveries among all declared positives. In cases where no features were declared significant (i.e., true positives = false positives = 0), we conservatively set the FDR to 0 to avoid undefined values.

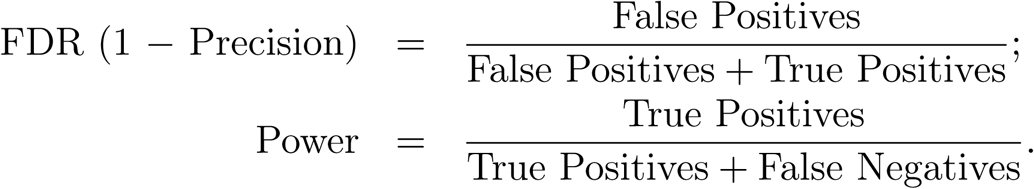

### Data Description

In this study, we analyzed two bulk RNA-Seq datasets, four single-cell RNA-Seq datasets, and one microbiome dataset, all obtained from publicly available sources as detailed below. Each dataset underwent initial filtering to remove low-quality features or samples, followed by normalization procedures tailored to its data type and context. Where feasible, we standardized preprocessing choices to ensure comparability while accommodating modality-specific requirements. For each dataset, we performed differential analysis between biologically meaningful case and control groups.

**LIHC (Bulk RNA-Seq):** A bulk RNA-Seq dataset consisting of 50 paired tumor (case) and adjacent normal (control) liver tissue samples. Sourced from the GDC Xena Hub^50^: https://xenabrowser.net/datapages/?hub=https://gdc.xenahubs.net:443,releasev18.0.
**GTEx (Bulk RNA-Seq):** A bulk RNA-Seq dataset from healthy human heart tissue, comparing the left ventricle (N = 386, case) with the atrial appendage (N = 372, control). Obtained from the GTEx Portal^59^: https://storage.googleapis.com/gtex_analysis_v8/rna_seq_data/GTEx_Analysis_2017-06-05_v8_RNASeQCv1.1.9_gene_reads.gct.gz.
**Brain (scRNA-Seq):** A non-UMI single-cell RNA-Seq dataset sourced from the SC2P R package ^102,103^, available via GEO accession GSE67835. The comparison was between astrocytes (N = 62) and oligodendrocytes (N = 38).
**PBMC (scRNA-Seq):** A single-cell RNA-Seq dataset of human immune cells from 10x Genomics ^111^, available at: https://support.10xgenomics.com/single-cell-gene-expression/datasets/1.1.0/pbmc3k. The analysis compared CD4^+^ T cells with CD8^+^ T cells.
**Kidney (scRNA-Seq):** A single-cell RNA-Seq dataset from GEO accession GSE107585 containing 43,745 cells and expression profiles for 16,273 genes. We analyzed a 10,000-cell subset from Wang *et al.* ^98^, comparing cells annotated as group C3 and C4.
**Petropoulos (scRNA-Seq):** A single-cell RNA-Seq dataset profiling human embryonic cells from embryonic day 3 (E3; N = 81) and embryonic day 4 (E4; N = 190). Data are available from ArrayExpress: https://www.ebi.ac.uk/biostudies/arrayexpress/studies/E-MTAB-3929 and mirrored at https://github.com/gongx030/scDatasets^77^.
**iHMP (Microbiome):** A microbiome dataset derived from the Integrative Human Microbiome Project (iHMP) ^57^, accessed in analysis-ready format from: https://github.com/WillNickols/maaslin3_benchmark. The comparison was between individuals with IBD (case) and non-IBD (control).

## Supplementary Materials for

### 1 Supplementary Table

**Table S1:**
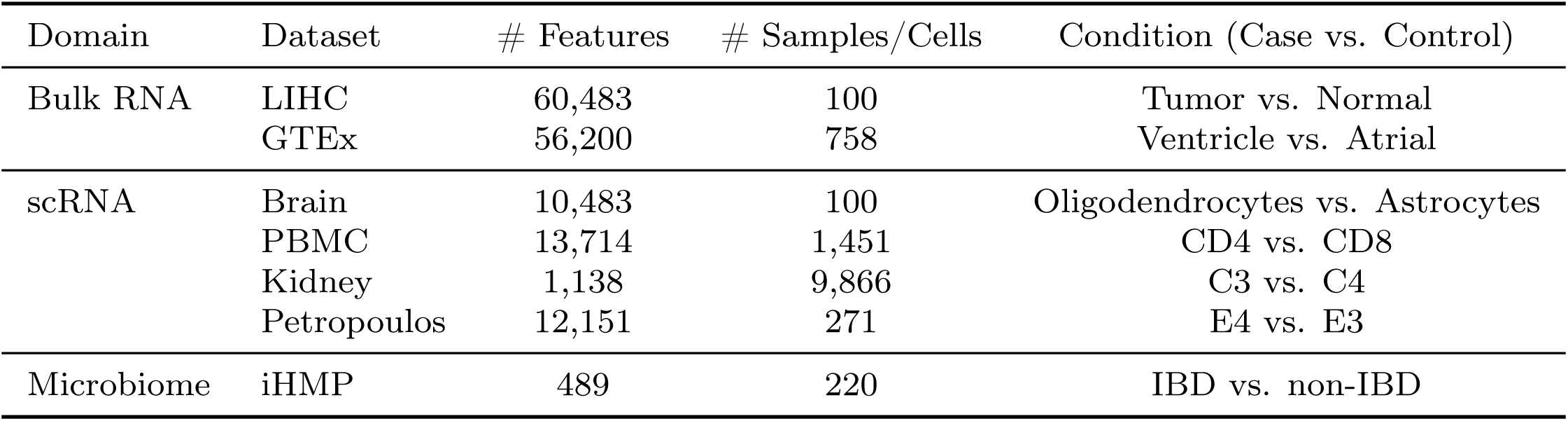
Summary of real datasets analyzed in the study. Overview of datasets from bulk RNA-Seq, scRNA-Seq, and microbiome domains, along with feature/sample counts and the condition labels used for differential analysis.

### 2 Supplementary Figures

**Figure S1:**
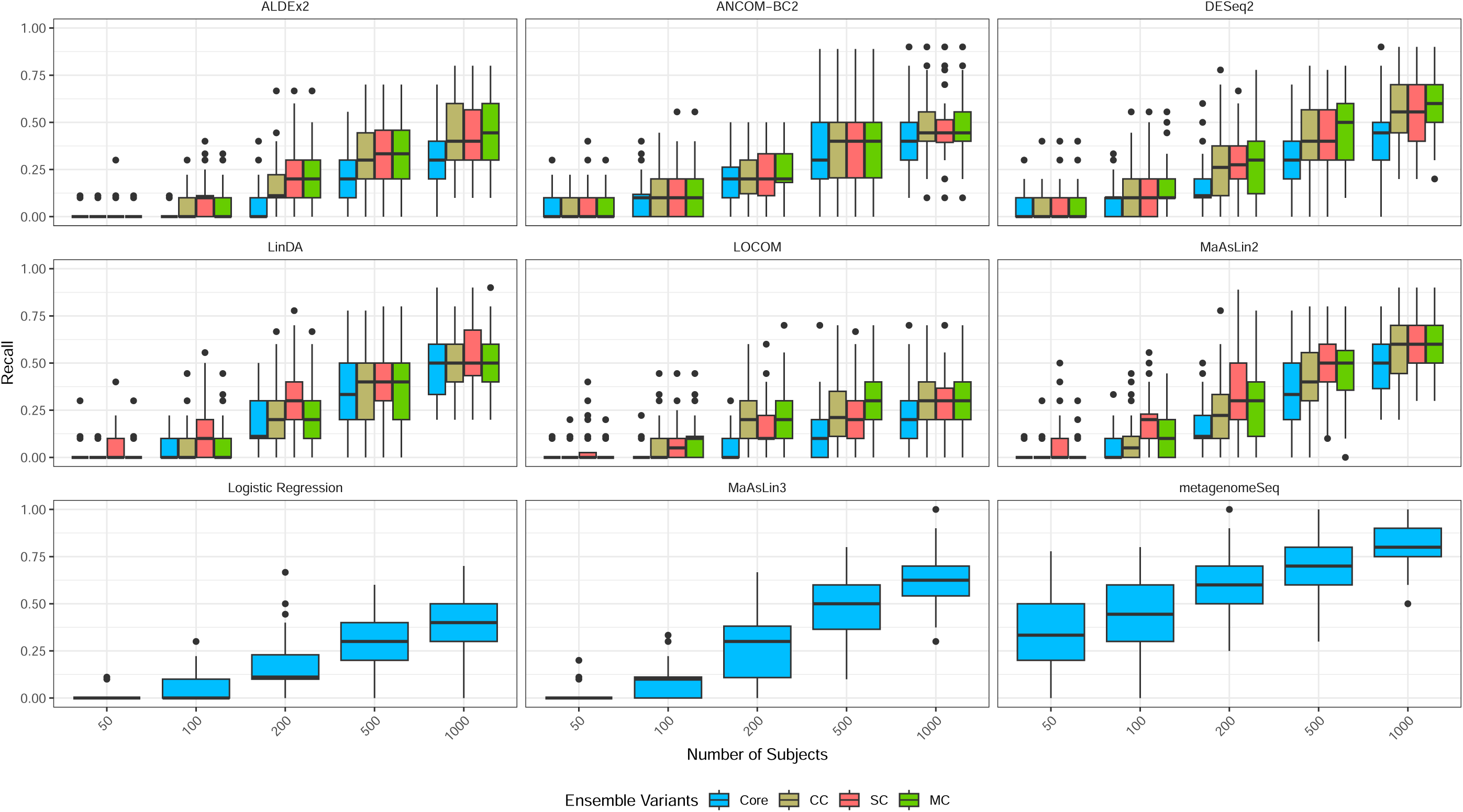
Recall performance of microbiome differential abundance methods and their LR-based ensembles under small effect sizes and varying sample sizes. Common microbiome DAA methods and their LR-based ensemble variants were evaluated on 100 synthetic datasets generated using SparseDOSSA2. Effect sizes were drawn uniformly from the range [1, 2], with the number of features fixed at 100. Recall values are summarized as boxplots across methods, with each point representing one simulated dataset. Higher recall indicates improved sensitivity in detecting true differential features.

**Figure S2:**
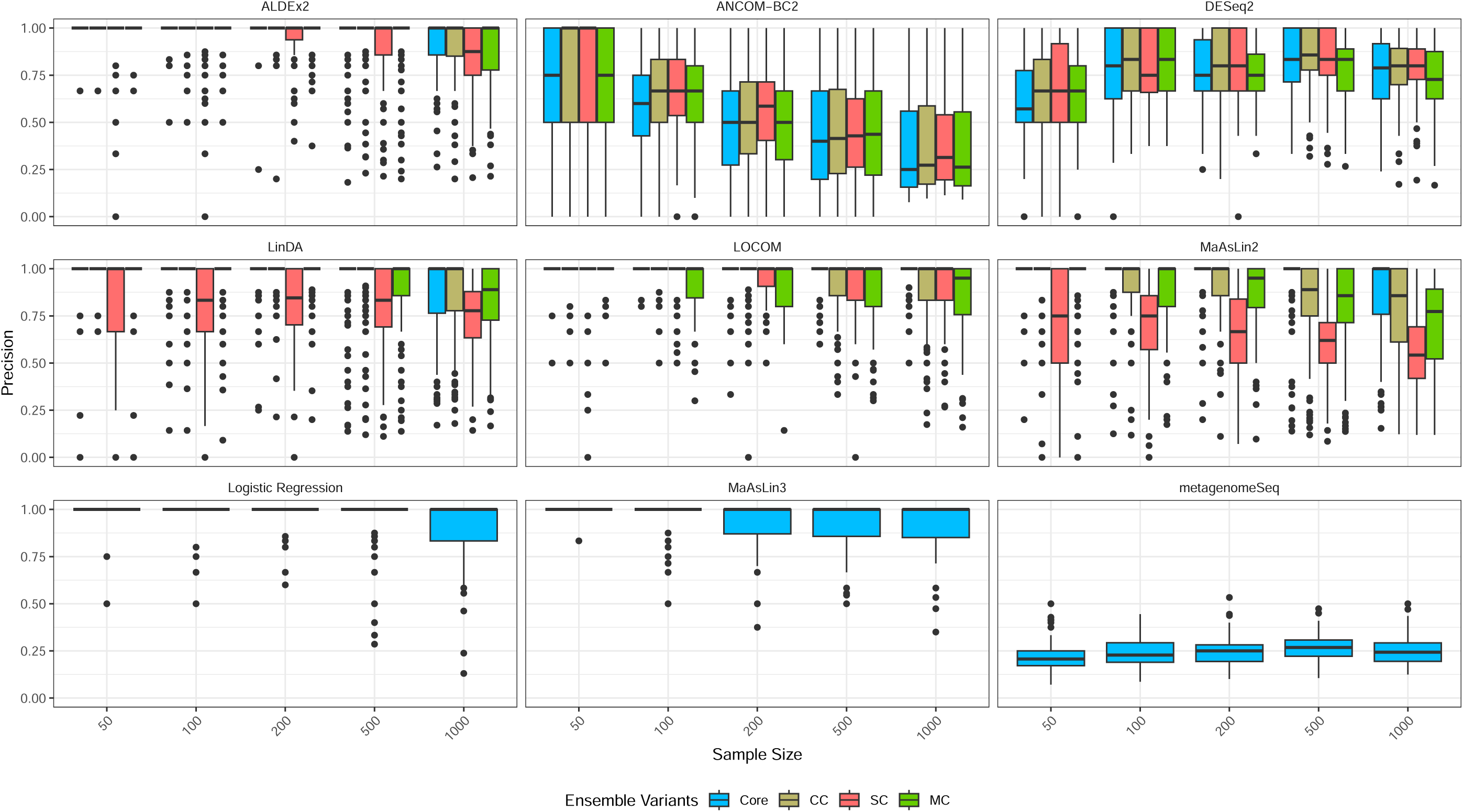
Precision performance of microbiome differential abundance methods and their LR-based ensembles under moderate effect sizes and varying sample sizes. Common microbiome DAA methods and their LR-based ensemble variants were evaluated on 100 synthetic datasets generated using SparseDOSSA2. Effect sizes were drawn uniformly from the range [2.5, 5], with the number of features fixed at 100. Precision values are summarized as boxplots across methods, with each point representing one simulated dataset. Higher precision indicates improved specificity in detecting true differential features.

**Figure S3:**
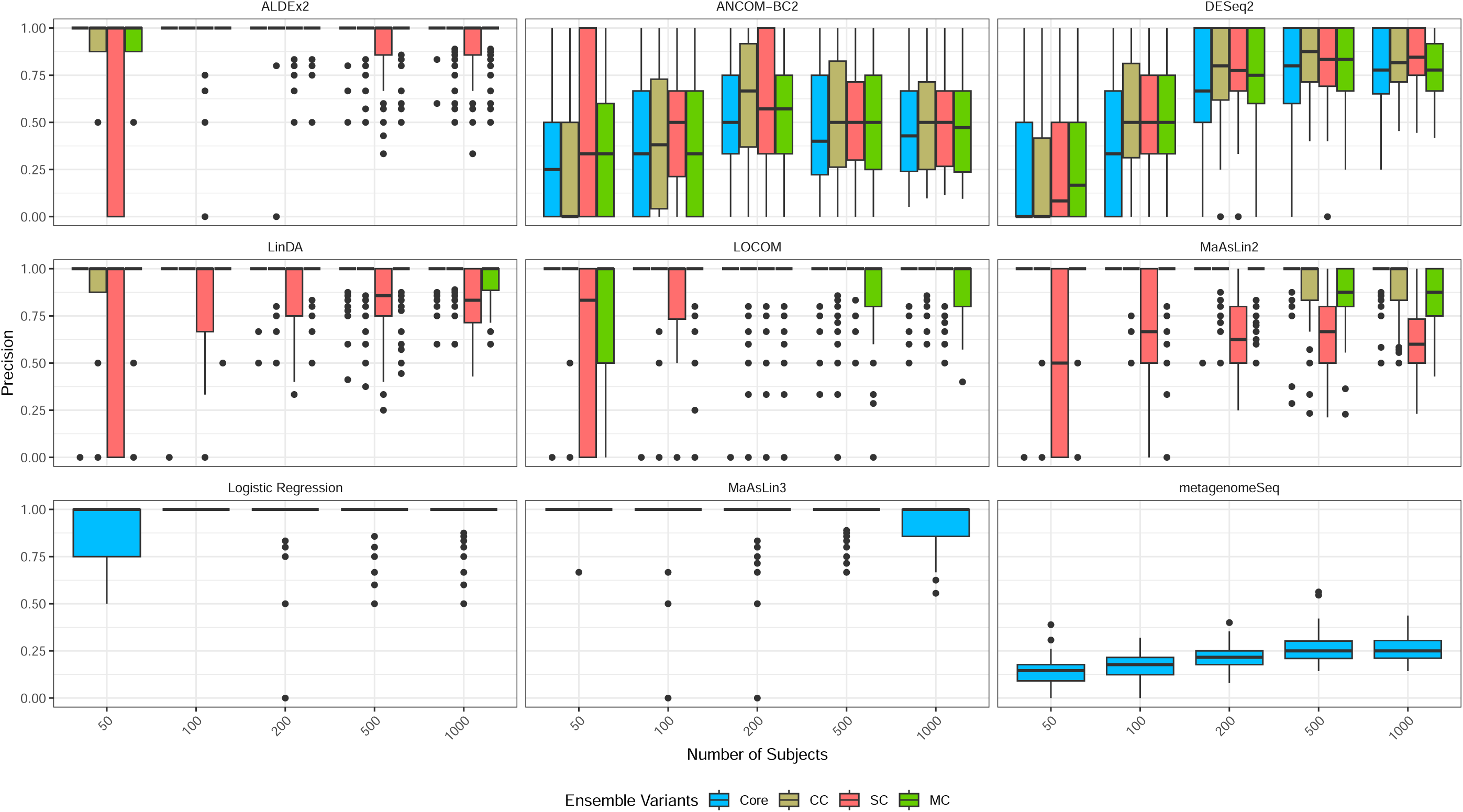
Precision performance of microbiome differential abundance methods and their LR-based ensembles under small effect sizes and varying sample sizes. Common microbiome DAA methods and their LR-based ensemble variants were evaluated on 100 synthetic datasets generated using SparseDOSSA2. Effect sizes were drawn uniformly from the range [1, 2], with the number of features fixed at 100. Precision values are summarized as boxplots across methods, with each point representing one simulated dataset. Higher precision indicates improved specificity in detecting true differential features.

**Figure S4:**
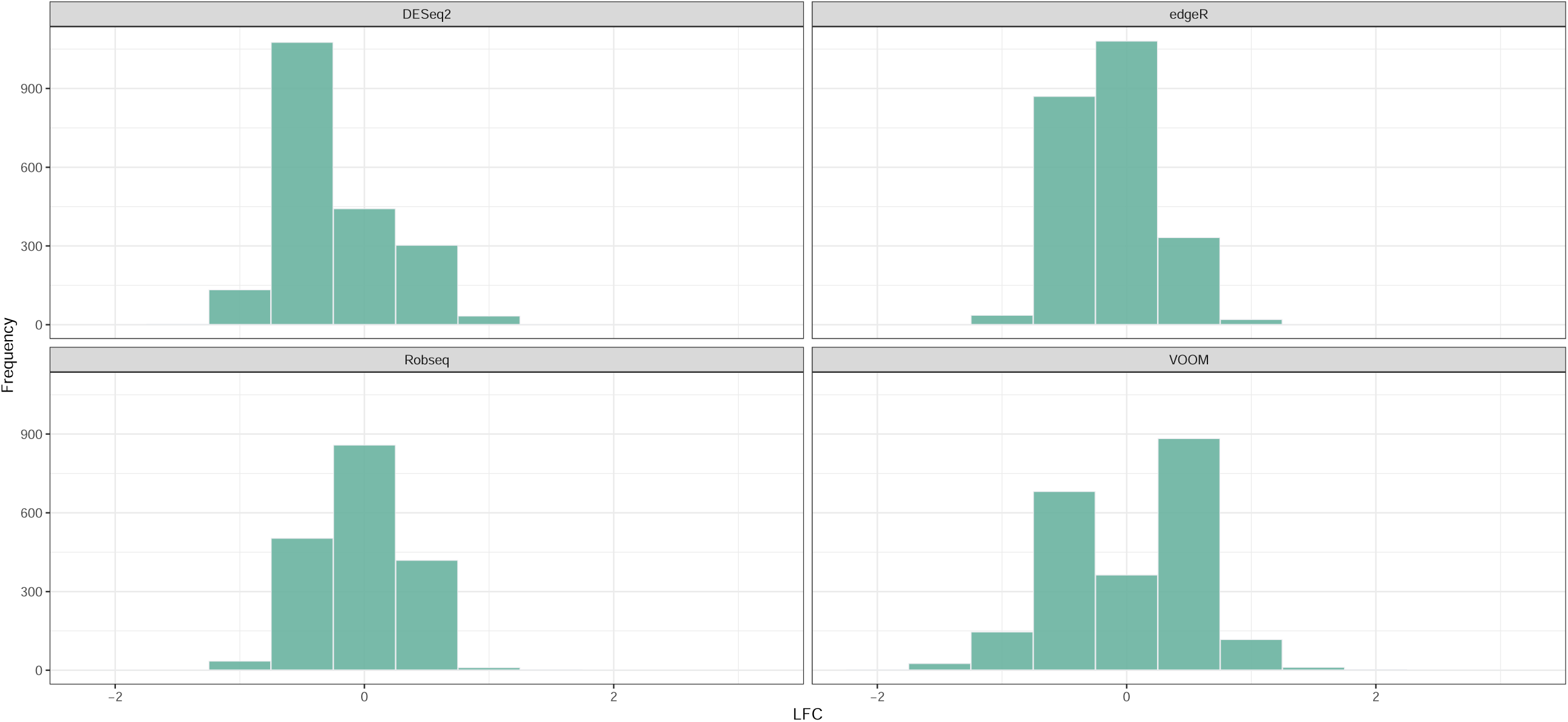
Histogram of effect sizes for all DE genes identified by *DAssemble* (SC variant) in the LIHC dataset. dearseq was excluded as it does not report effect sizes.

**Figure S5:**
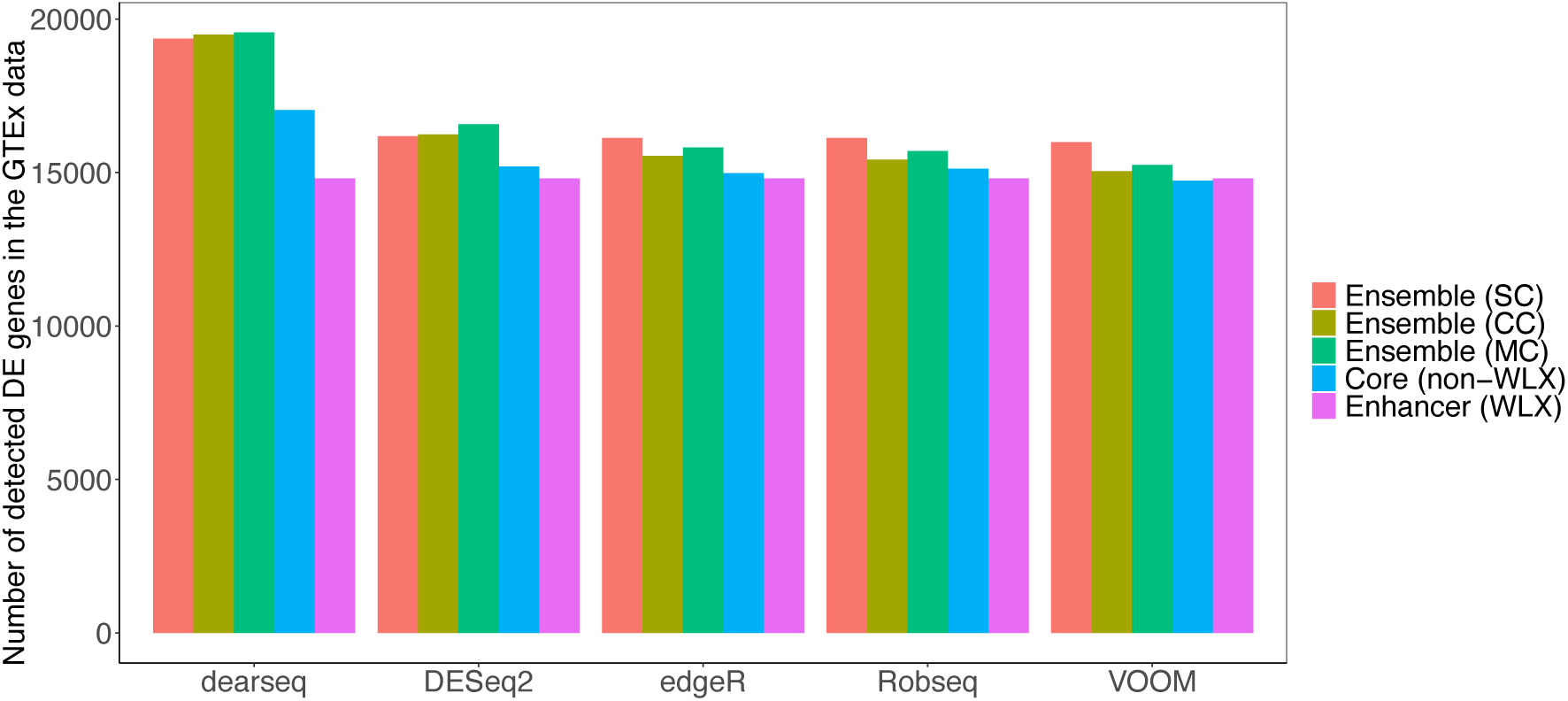
Summary of DE genes detected by various bulk RNA-Seq differential expression models on the GTEx dataset^1^. The analysis includes a WLX enhancer model, five non-WLX core models (dearseq, DESeq2, edgeR, RobSeq, and VOOM), and their ensemble variants constructed using CC, SC, and MC. In each case, at least one ensemble method identified a significantly greater number of DE genes (FDR < *α* = 0.05) compared to its corresponding non-ensemble counterpart, demonstrating the enhanced sensitivity of the ensemble approaches in capturing biological signals in this dataset.

**Figure S6:**
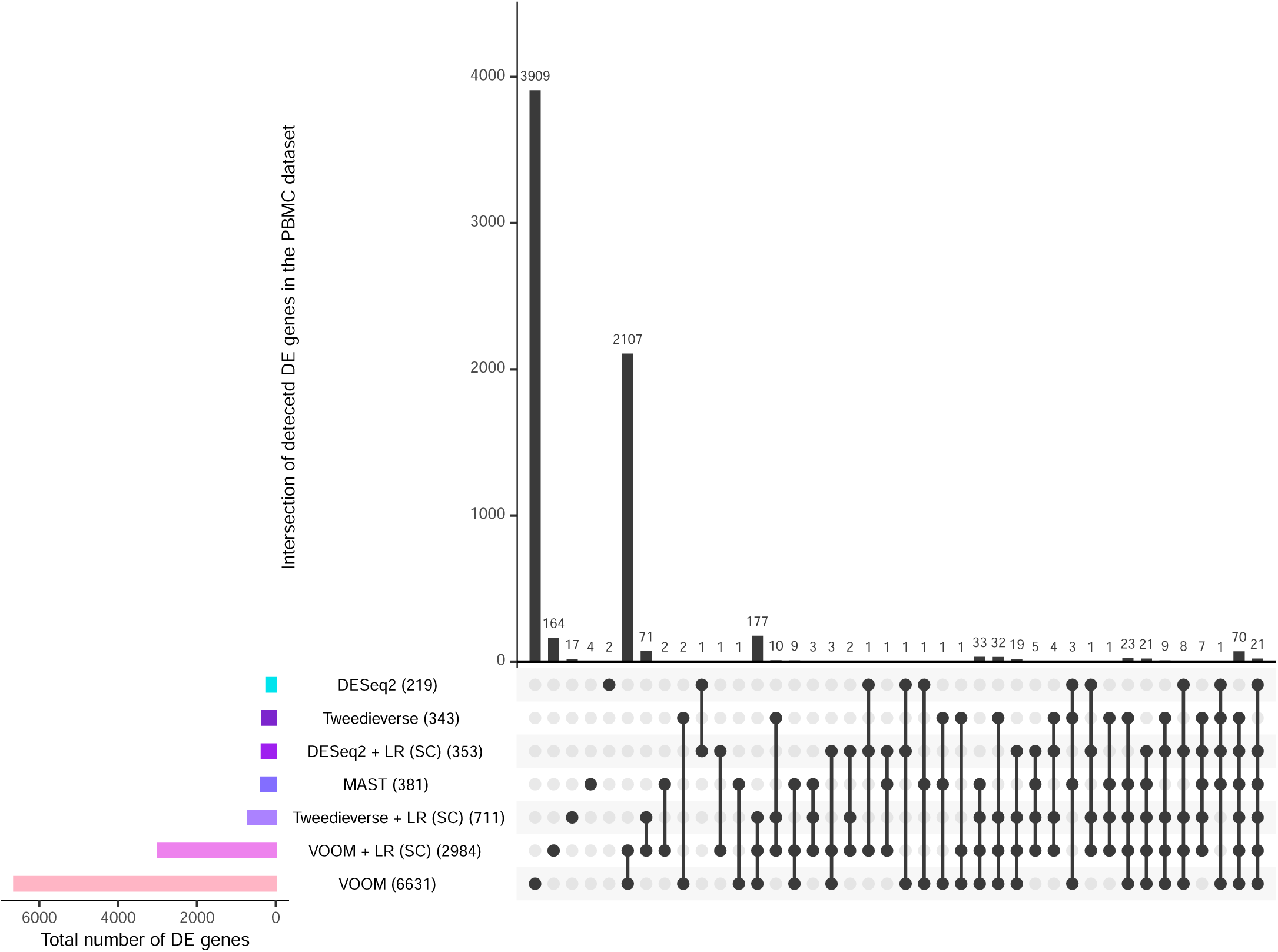
Overlap of DE genes detected by five scRNA-Seq methods in the PBMC data. Five scRNA-Seq differential expression methods were applied to identify DE genes between CD8^+^ (*N* = 271) and CD4^+^ T cells (*N* = 1180) in the PBMC dataset ^5^, across a total of 13,714 genes. Genes with observed FDR below *α* = 0.05 were considered DE. Numbers in parentheses indicate the total number of DE genes detected by each method. The results illustrate both shared and method-specific gene-level discoveries.

**Figure S7:**
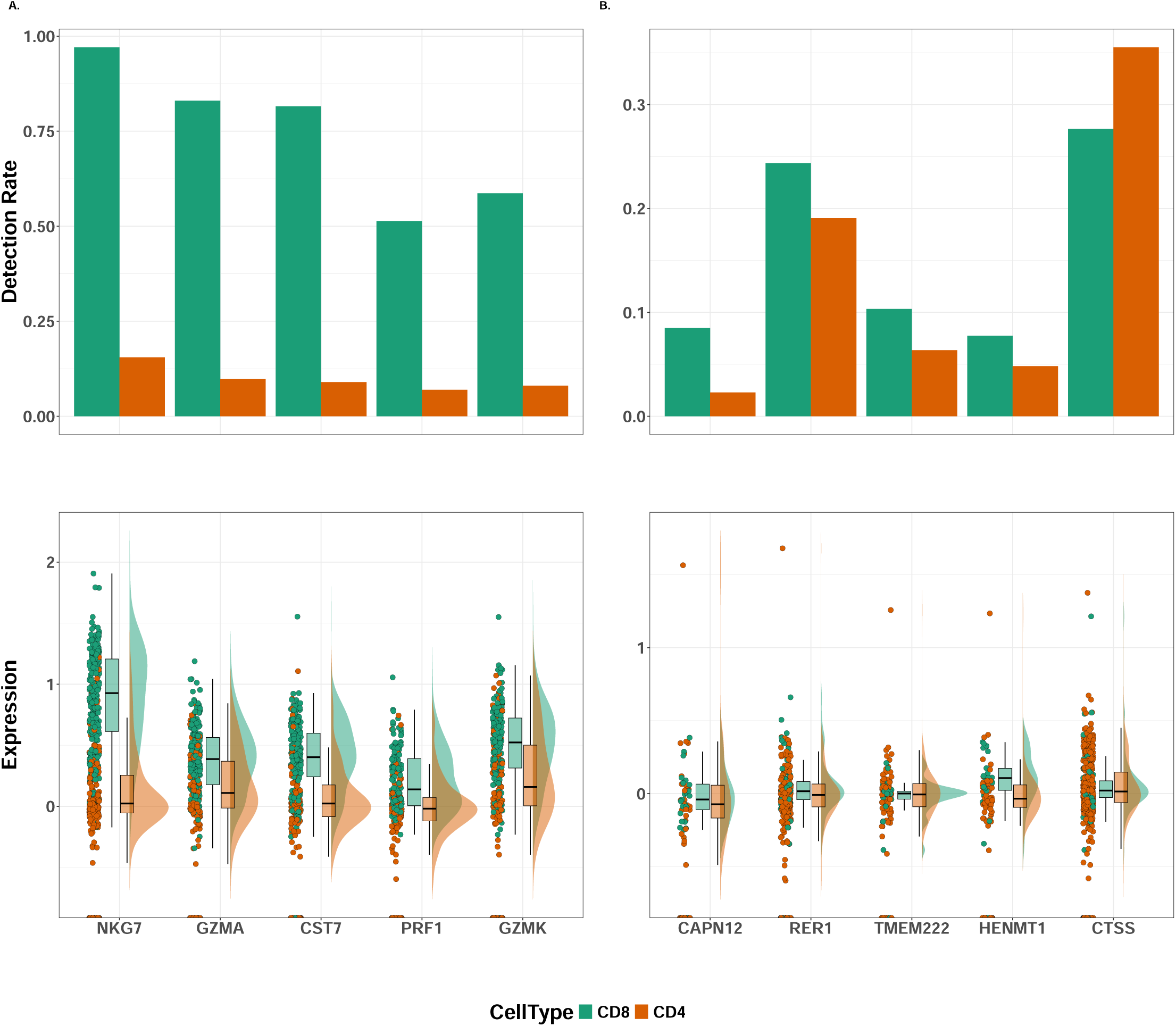
Raincloud plots of the top 5 DE genes in the PBMC dataset. Genes were selected based on adjusted p-values below *α* = 0.05. Bar charts represent the prevalence of each gene, while violin plots, overlaid with box plots and scatter points, illustrate their distribution and abundance across the two cell types. Panel **A** includes genes for which both components of the two-part model yielded adjusted p-values below 0.05, whereas panel **B** includes genes with adjusted p-values greater than 0.5 in both components.

**Figure S8:**
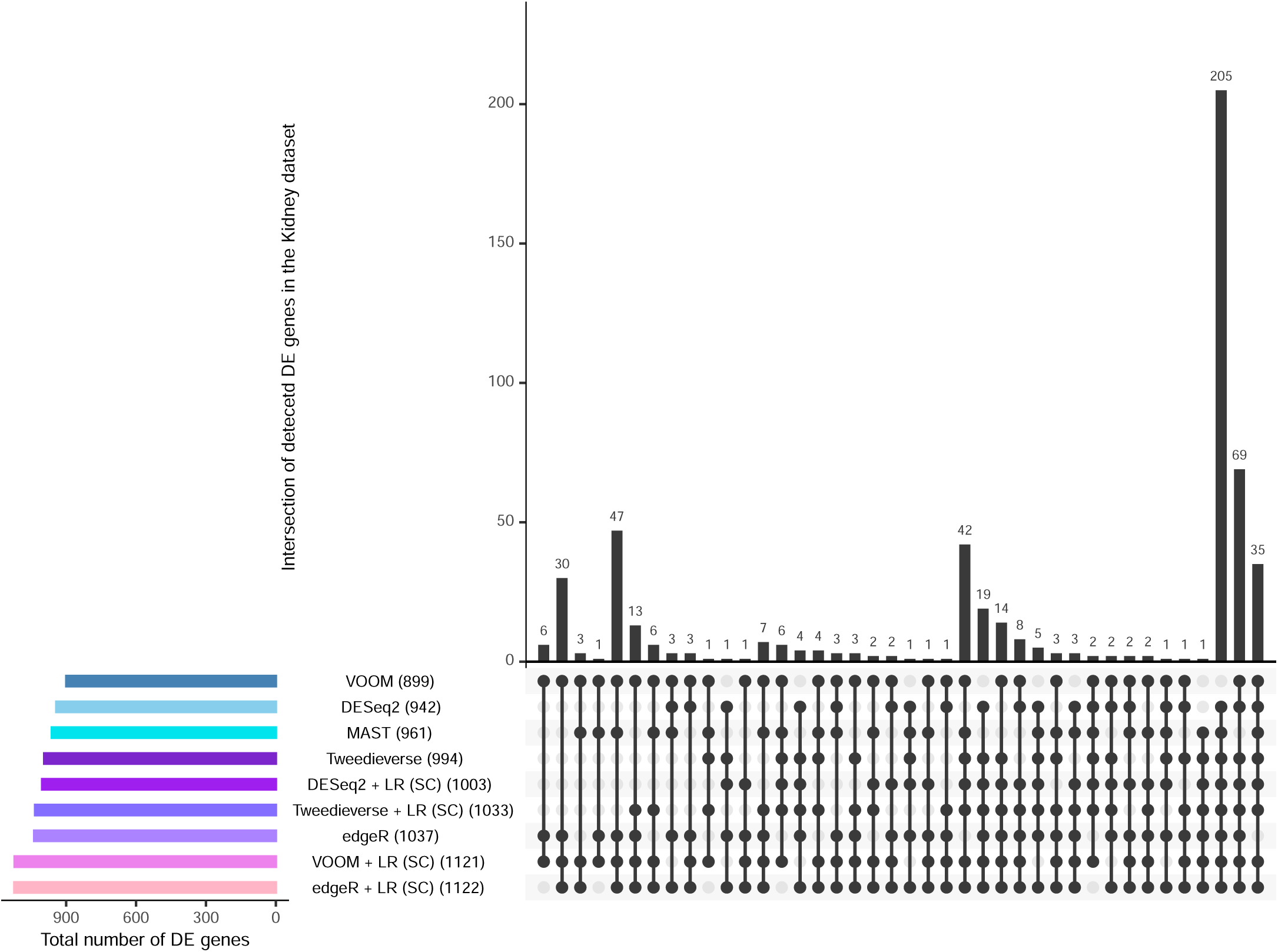
Overlap of DE genes detected by five scRNA-Seq methods in the Kidney data. Using five scRNA-Seq differential expression methods, the number of DE genes (out of 1138) detected between C3 (epithelial cells) (*N* = 9, 129 cells) versus C4 (immune cells) (*N* = 737 cells) clusters. Genes with observed FDR below *α* = 0.05 were considered DE. Numbers in parentheses indicate the total number of DE genes detected by each method. The results illustrate both shared and method-specific gene-level discoveries.

**Figure S9:**
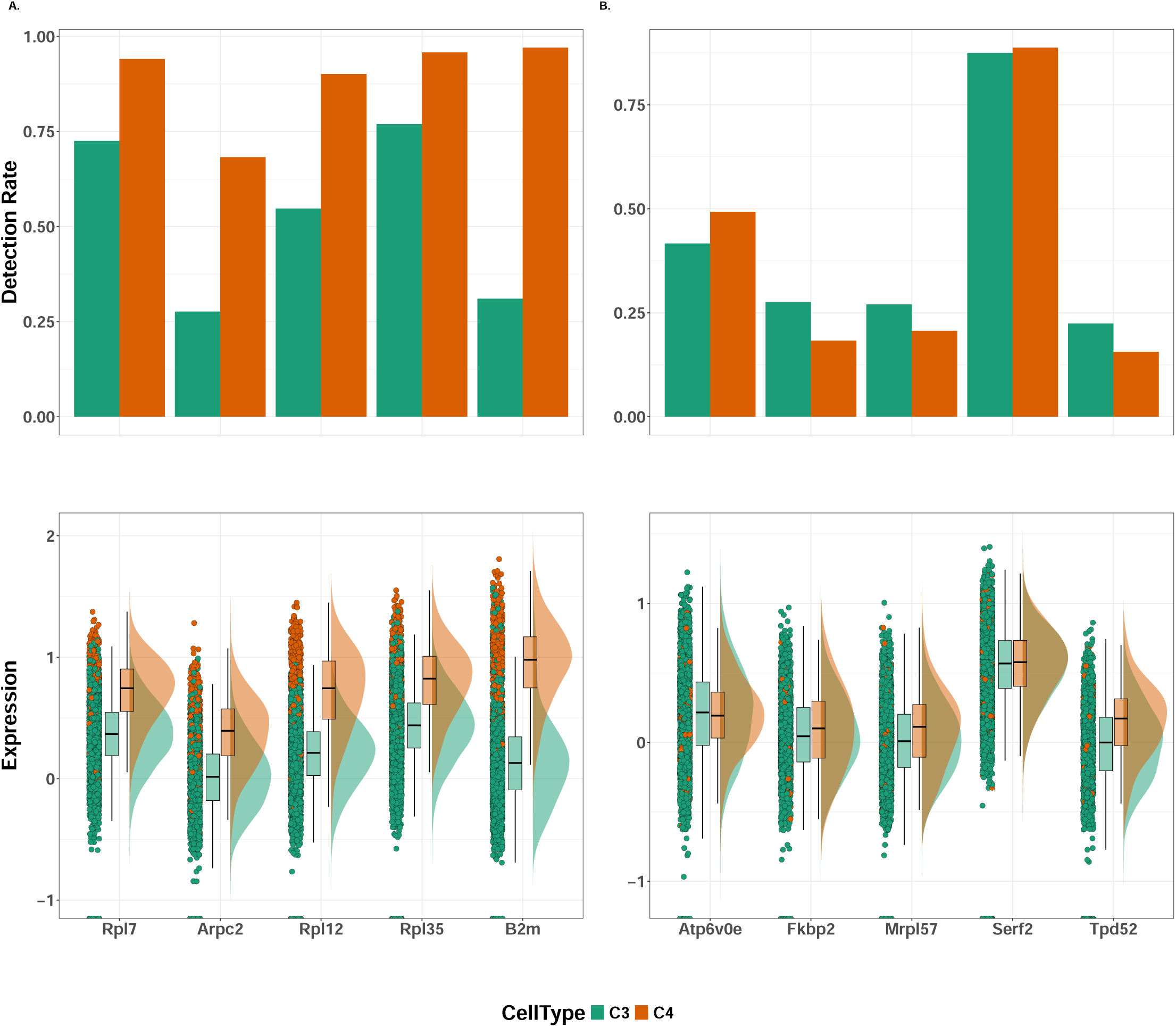
Raincloud plots of the top 5 DE genes in the Kidney dataset. Genes were selected based on adjusted p-values below *α* = 0.05. Bar charts represent the prevalence of each gene, while violin plots, overlaid with box plots and scatter points, illustrate their distribution and abundance across the two cell types. Panel **A** includes genes for which both components of the two-part model yielded adjusted p-values below 0.05, whereas panel **B** includes genes with adjusted p-values greater than 0.5 in both components.

**Figure S10:**
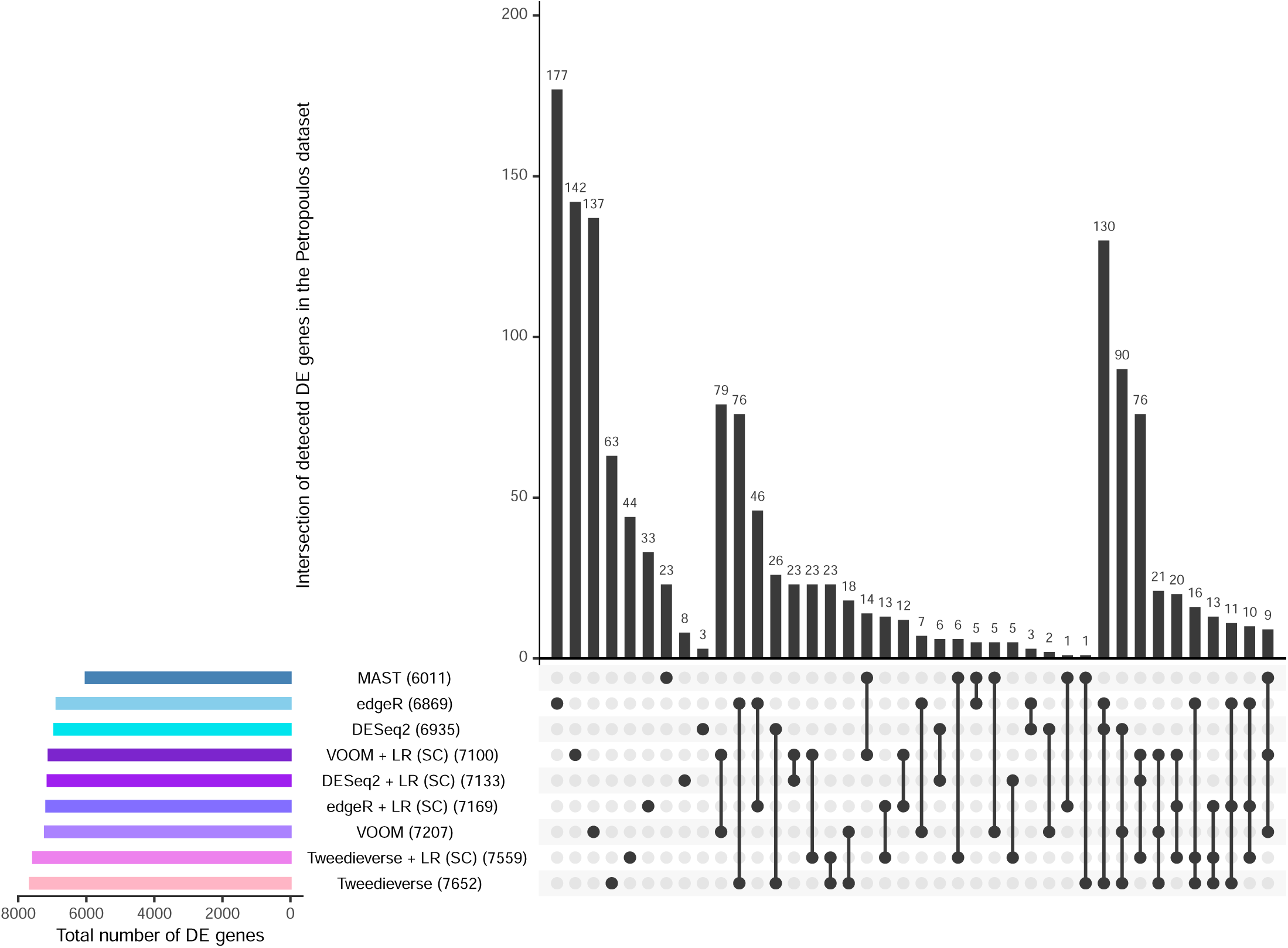
Overlap of DE genes detected by five scRNA-Seq methods in the Petropoulos dataset^3^. Five scRNA-Seq differential expression methods were applied to identify DE genes between embryonic day 3 (E3, *N* = 81) and embryonic day 4 (E4, *N* = 190) groups. This dataset was previously analyzed by Miao *et al.* ^2^. Following the gene-filtering strategy of Sekula *et al.* ^4^, genes expressed in fewer than 20% of cells were excluded, resulting in a final set of 12,151 genes. Genes with observed FDR below *α* = 0.05 were considered DE. Numbers in parentheses indicate the total number of DE genes detected by each method.

**Figure S11:**
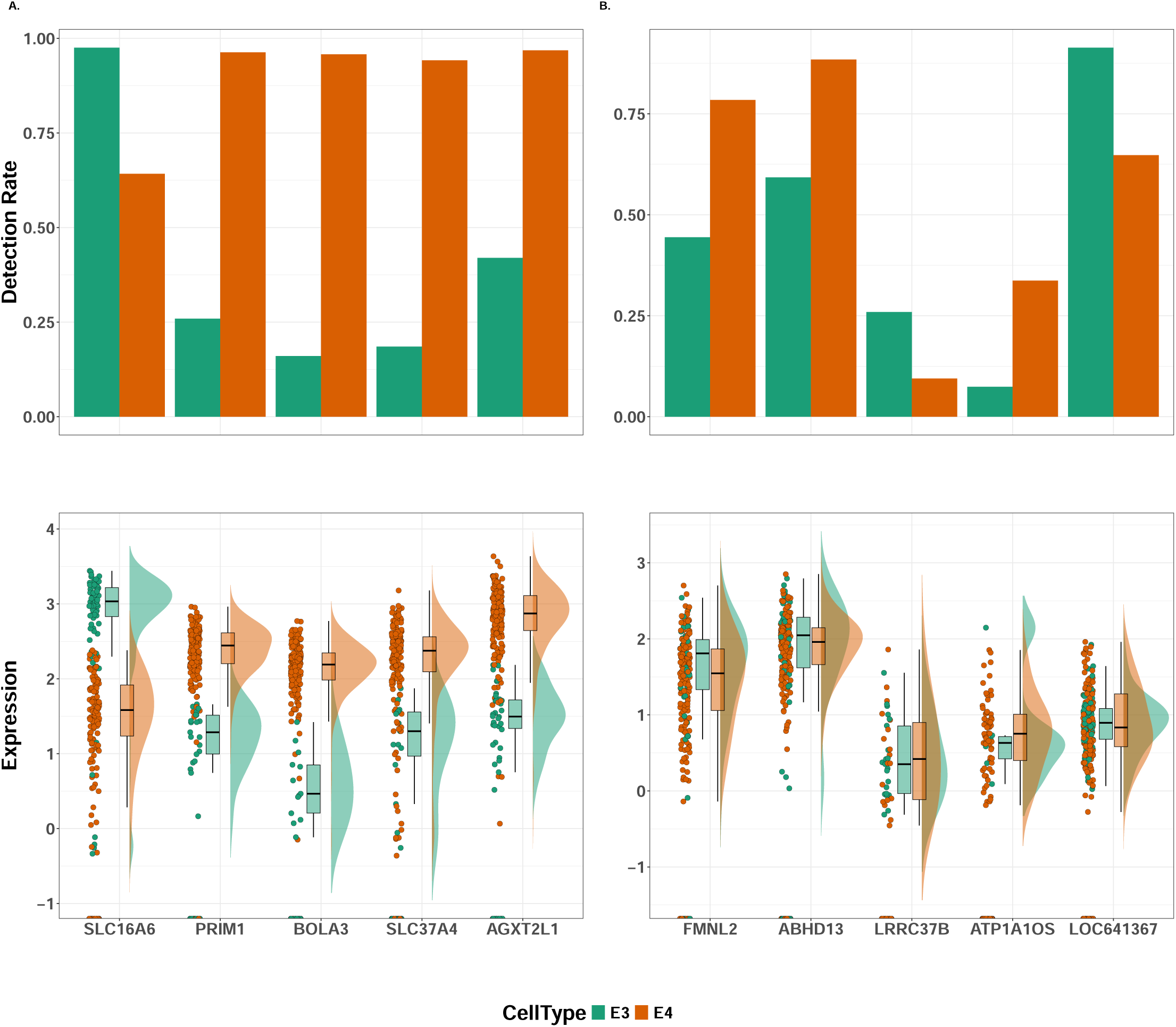
Raincloud plots of top 5 DE genes identified in the Petropoulos dataset^3^. Genes were selected based on adjusted p-values below *α* = 0.05. Bar charts represent the prevalence of each gene, while the violin plots, overlaid with box plots and scatter points, illustrate their distribution and abundance across the two cell types. Panel **A** includes genes for which both components of the two-part model yielded adjusted p-values below 0.05, whereas panel **B** includes genes with adjusted p-values greater than 0.5 in both components.

**Figure S12:**
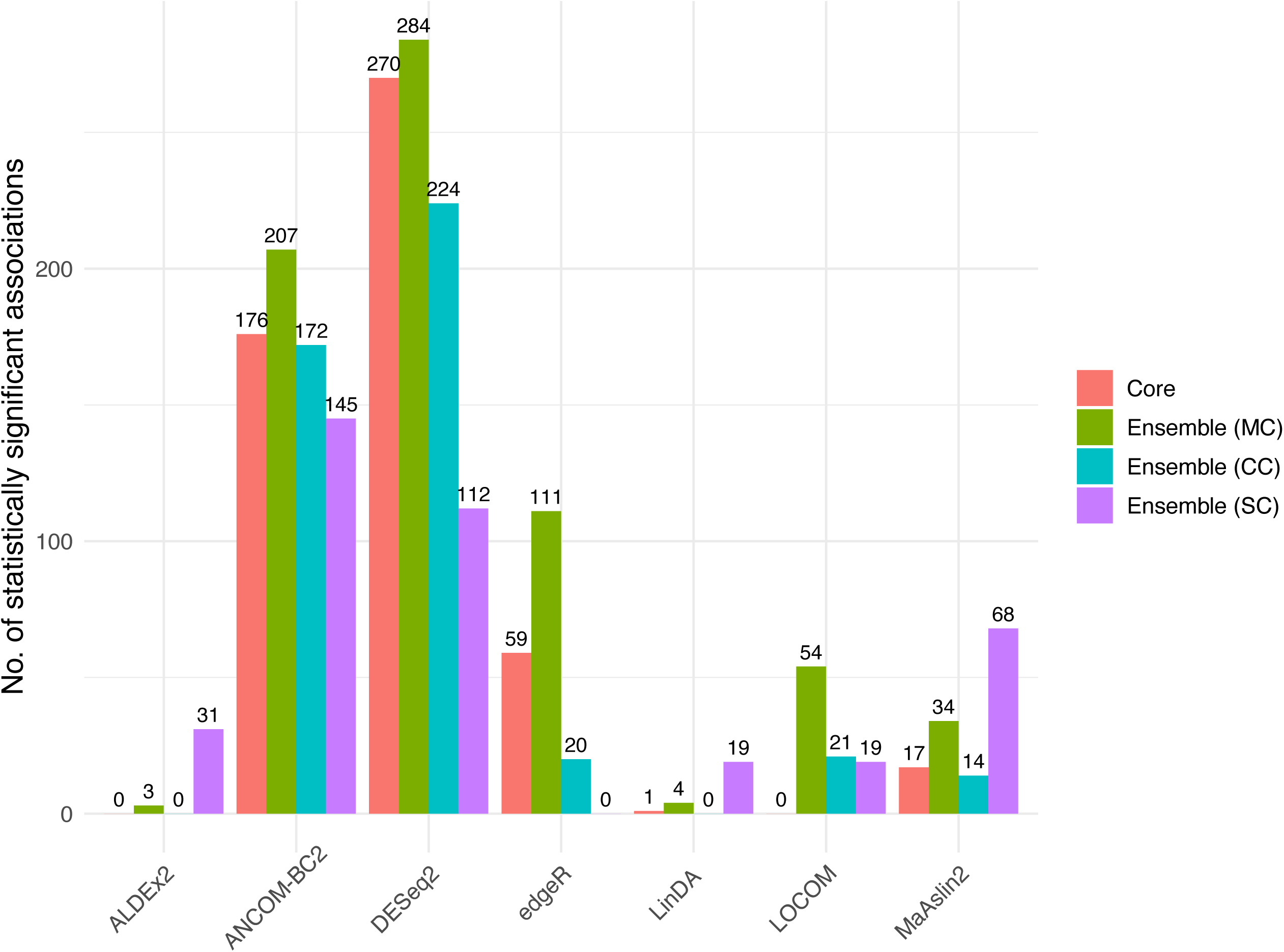
Statistically significant associations identified by LR-enhanced versions of seven core microbiome DAA methods (ALDEx2, ANCOM-BC2, DESeq2, edgeR, LinDA, LOCOM, and MaAsLin2) across all ensemble variants in the iHMP dataset. Results are grouped by core method (x-axis) and colored by ensemble variant (MC, SC, CC).

**Figure S13:**
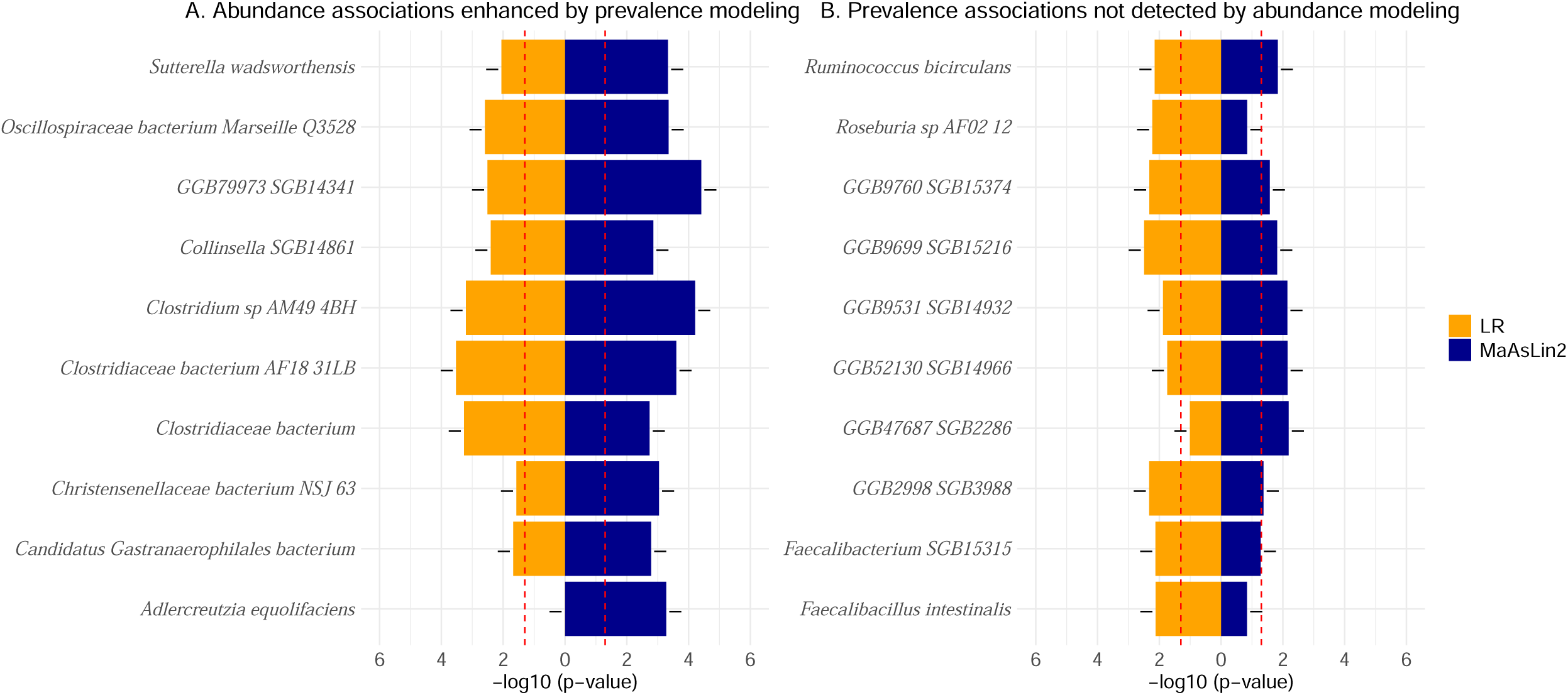
Top microbial species associated with IBD identified in the iHMP dataset using the MC ensemble approach. Using MaAsLin2 and LR as the base and enhancer models, respectively, we report the top 10 microbial species ranked by adjusted p-values from the MC ensemble, which integrates p-values from MaAsLin2 and LR before FDR correction. The left panel presents *−* log_10_(*p*) values from MaAsLin2 and LR for species identified as significant by both methods. The right panel displays species uniquely detected by the ensemble, despite receiving low ranks from MaAsLin2 alone. Positive (+) and negative (–) signs denote the direction of association with IBD. A red dotted line in these plots represents the nominal type-I error rate, set at 0.05.

**Figure S14:**
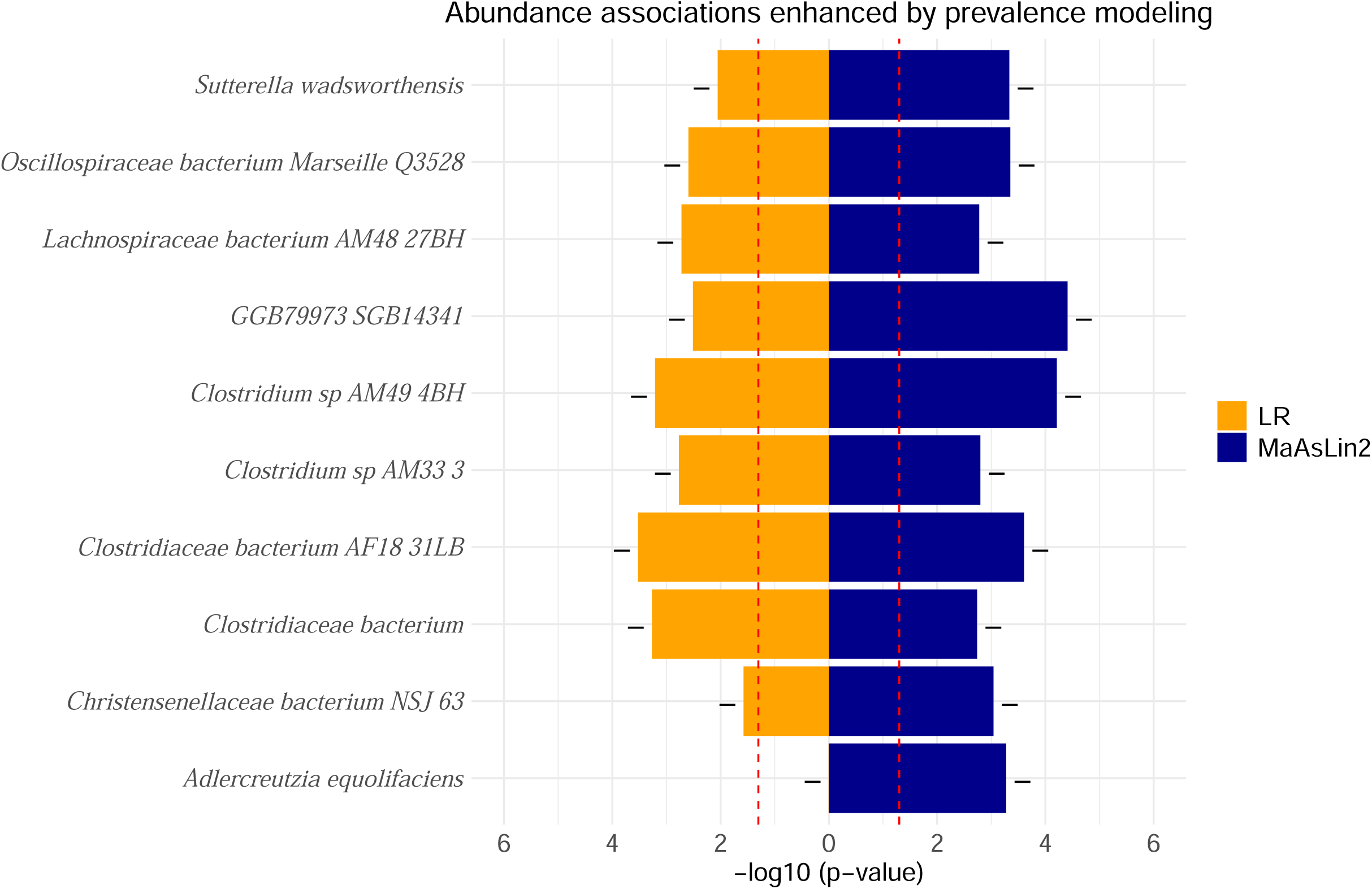
Top microbial species associated with IBD identified in the iHMP dataset using the CC ensemble approach. Using MaAsLin2 and LR as the base and enhancer models, respectively, we report the top 10 microbial species ranked by adjusted p-values from the CC ensemble, which integrates p-values from MaAsLin2 and LR before FDR correction. Only microbial species identified as significant by both methods are shown, as no species were uniquely detected by the CC ensemble. Plus (+) and minus (–) signs indicate the direction of association with IBD. A red line in these plots represents the nominal type-I error rate, set at 0.05.

### 3 Supplementary Data

**Supplementary Datasets S1–S7: Full results from differential association analyses across datasets**. Each file lists statistically significant associations (FDR < 0.05) between metadata groups. **S1:** LIHC (Tumor vs. Normal), **S2:** GTEx (Atrium vs. Ventricle), **S3:** Brain (Oligodendrocytes vs. Astrocytes), **S4:** PBMC (CD4 vs. CD8), **S5:** Kidney (C3 vs. C4), **S6:** Petropoulos (E3 vs. E4), and **S7:** iHMP (IBD vs. non-IBD). Features are sorted by minimum FDR-adjusted p-values. For each feature, the coefficient estimates, standard errors, and associated two-tailed p-values are reported.

**Supplementary Dataset S8:** Results showing the overlap of DE genes identified across all core models and SC ensemble variants in the LIHC dataset. Notably, the ensemble variants uniquely detected thousands of DE genes not identified by the core models alone, with varying degrees of overlap observed among the ensemble approaches.

**Supplementary Dataset S9:** In the LIHC dataset, the ensemble variants uncovered DE genes with subtle effect sizes that were missed by the core models, highlighting the enhanced sensitivity of the ensemble approach. DE genes consistently detected across ensemble strategies (CC, MC, SC) were compiled and summarized.

**Supplementary Dataset S10: Biological relevance of the top genes uniquely identified by *DAssemble* in the LIHC dataset.** DE genes consistently detected across ensemble strategies (CC, MC, SC) were compiled, and the top 10 were selected based on FDR-adjusted p-values.

